# Satellite DNA landscapes after allotetraploidisation of quinoa (*Chenopodium quinoa*) reveal unique A and B subgenomes

**DOI:** 10.1101/774828

**Authors:** Tony Heitkam, Beatrice Weber, Ines Walter, Charlotte Ost, Thomas Schmidt

## Abstract

If two related plant species hybridise, their genomes are combined within a single nucleus, thereby forming an allotetraploid. How the emerging plant balances two co-evolved genomes is still a matter of ongoing research. Here, we focus on satellite DNA (satDNA), the fastest turn-over sequence class in eukaryotes, aiming to trace its emergence, amplification and loss during plant speciation and allopolyploidisation. As a model, we used *Chenopodium quinoa* Willd. (quinoa), an allopolyploid crop with 2n=4x=36 chromosomes. Quinoa originated by hybridisation of an unknown female American *Chenopodium* diploid (AA genome) with an unknown male Old World diploid species (BB genome), dating back 3.3 to 6.3 million years. Applying short read clustering to quinoa (AABB), *C. pallidicaule* (AA), and *C. suecicum* (BB) whole genome shotgun sequences, we classified their repetitive fractions, and identified and characterised seven satDNA families, together with the 5S rDNA model repeat. We show unequal satDNA amplification (two families) and exclusive occurrence (four families) in the AA and BB diploids by read mapping as well as Southern, genomic and fluorescent *in situ* hybridisation. As *C. pallidicaule* harbours a unique satDNA profile, we are able to exclude it as quinoa’s parental species. Using quinoa long reads and scaffolds, we detected only limited evidence of interlocus homogenisation of satDNA after allopolyploidisation, but were able to exclude dispersal of 5S rRNA genes between subgenomes. Our results exemplify the complex route of tandem repeat evolution through *Chenopodium* speciation and allopolyploidisation, and may provide sequence targets for the identification of quinoa’s progenitors.

## INTRODUCTION

Quinoa (*Chenopodium quinoa* Willd.) is an allotetraploid crop, domesticated in the South-American Andes for at least 8000 years (Dillehay et al., 2007). Due to the ability to grow on marginal soils in short vegetation periods, the high tolerance for abiotic stresses such as cold and UV radiation, and the high nutritional value of its grains, quinoa is ranked as a “high potential” crop with priority for sustainable agriculture (www.fao.org, Zurita-Silva et al., 2014). Having an allotetraploid origin with 2n=4x=36 chromosomes, quinoa was derived from hybridisation of a female American *Chenopodium* diploid (A genome) with a male Old World diploid (B genome), dating back 3.3 to 6.3 million years (Kolano *et al*., 2012, Štorchová *et al*., 2015, Walsh *et al*., 2015, Jarvis *et al*., 2017, Maughan *et al*., 2019). Potential diploid progenitors with B genomes have approximately 30 % larger genomes than A genome species (approximately 0.6 and 0.9 Gb; Kolano et al., 2016). Thus, for the quinoa genome size of 1.45 to 1.5 Gb (Palomino et al., 2008), additivity of A and B genomes without genome up- or downsizing was postulated (Kolano et al., 2016). With ongoing sequencing efforts, three independent quinoa reference genomes have been generated (Yasui et al., 2016, Jarvis et al., 2017, Zou et al., 2017).

Phylogenetically, quinoa is a member of the Amaranthaceae (formerly Chenopodiaceae), which encompasses many crops, some of them with genome sequence data available, e.g. *Beta vulgaris* (sugar beet, Dohm et al., 2014)*, Spinacia oleracea* (spinach, Xu et al., 2017), and *Amaranthus hypochondriacus* (amaranth, Clouse et al., 2016). Quinoa belongs to the subfamily *Chenopodioideae*, which has recently been revised and split into multiple smaller genera such as *Chenopodiastrum*, *Oxybasis*, *Lipandra*, and *Dysphania*, in addition to the previously recognised genera *Chenopodium*, *Atriplex*, *Blitum*, and *Spinacia* (Fuentes-Bazan et al., 2012a, Fuentes-Bazan et al., 2012b).

To assign specific chromosomes of hybrids or polyploids to their respective parental genomes as well as to physically map DNA sequences along chromosomes, fluorescent and genomic *in situ* hybridisation (FISH and GISH) are valuable technologies (Schwarzacher et al., 1989, D’Hont, 2005, Chester et al., 2010, Mandáková et al., 2013, Jiang, 2019). In many *Chenopodium* species, the 5S- and 18S-5.8S-25S rRNA genes have already been sequenced and localized on chromosomes, including potential A and B genome progenitors of quinoa (Maughan et al., 2006, Kolano et al., 2012, Kolano et al., 2016, Kolano et al., 2019).

Fast-evolving tandem repeats such as satellite DNA (satDNAs) can serve as chromosomal landmarks and are often specific for individual chromosomes or subgenomes (Shapiro and von Sternberg, 2005, Heslop-Harrison and Schwarzacher, 2011, Schmidt *et al*., 2019). SatDNAs are non-translated and tandemly organised repeated units, often uniform within large chromosomal arrays. Maintenance of genome integrity, epigenetic regulation of gene expression, and centromere formation are among the main functional roles of satDNA (Melters *et al*., 2013, Zakrzewski *et al*., 2013, Zhang *et al*., 2013, Jagannathan *et al*., 2018).

On an evolutionary timescale, tandem repeats change rapidly and are among the first sequence classes to diversify in emerging species (Charlesworth *et al*., 1994, Oliver *et al*., 2013, Zhang *et al*., 2015, McCann *et al*., 2018, Bracewell *et al*., 2019). Accumulation of mutations (single nucleotide changes, indels) leads to the emergence of new satDNAs variants, which may spread and displace existing variants and often form homogenised arrays (Plohl et al., 2012, Garrido-Ramos, 2015). Allopolyploidisation of related species combines diverged repetitive DNA families (Koukalova et al., 2010, Vicient and Casacuberta, 2017), with a range of largely unpredictable consequences: In the new allopolyploid, repeats may be redistributed, reduced, replaced, newly combined, or selectively amplified. In quinoa, already some repetitive sequences have been identified, including also a satDNA family (Kolano *et al*., 2011, Orzechowska *et al*., 2018). However, genome wide satDNA profiles of quinoa and potential parental species are still needed to deduce the effects of speciation and allopolyploidisation on *Chenopodium* repeat evolution.

Here, we analysed the influence of speciation and polyploidisation on the tandem repeat landscape within *Chenopodium*. Next-generation genome sequences of allotetraploid *C. quinoa* and two putative diploid progenitors, *C. pallidicaule* (A genome) and *C. suecicum* (B genome) were subjected to comparative read clustering. We identified the major tandem repeats, including satDNAs and 5S rRNA genes, and examined their amplification in A and B genomes. Their higher order organisation and the interlocus homogenisation of A- and B-specific variants was investigated using *C. quinoa* long single molecule real-time (SMRT) reads. Combining GISH and FISH, we show that the satellite DNAs target A- and B-specific chromosomes in *C. quinoa* genomes and reveal a surprisingly low rate of inter-subgenome dispersion.

## RESULTS

### Identification of tandem repeats in *Chenopodium* genomes

In order to measure the contributions of the A and B subgenomes on the *C. quinoa* repeat composition, we compared the repeat fractions of *C. pallidicaule* (AA diploid), *C. suecicum* (BB diploid), and *C. quinoa* (AABB tetraploid) by three-way read clustering with equal amounts of shot gun Illumina reads. We used approximately 127.8 Mb of each genome as input for *RepeatExplorer*, enabling a representative read cluster analysis (Novák et al., 2013, Goubert et al., 2015, Weiss-Schneeweiss et al., 2015).

Based on the data we estimate the fraction of repetitive DNA to be 76.9 % for *C. pallidicaule*, 78.1 % for *C. suecicum*, and 73.2 % for *C. quinoa*. However, as diverged repeats escape the clustering threshold of 90 %, the real repeat content is likely higher. To quantify highly repetitive sequences, we comparatively analysed the largest 150 clusters (Figure 1) and, if possible, assigned the underlying sequences to Ty1-*copia* and Ty3-*gypsy* long terminal repeat (LTR) retrotransposons, pararetroviruses, long interspersed nuclear elements (LINEs), DNA transposons, rDNA, satDNA, and organellar DNA.

**Fig. 1:**
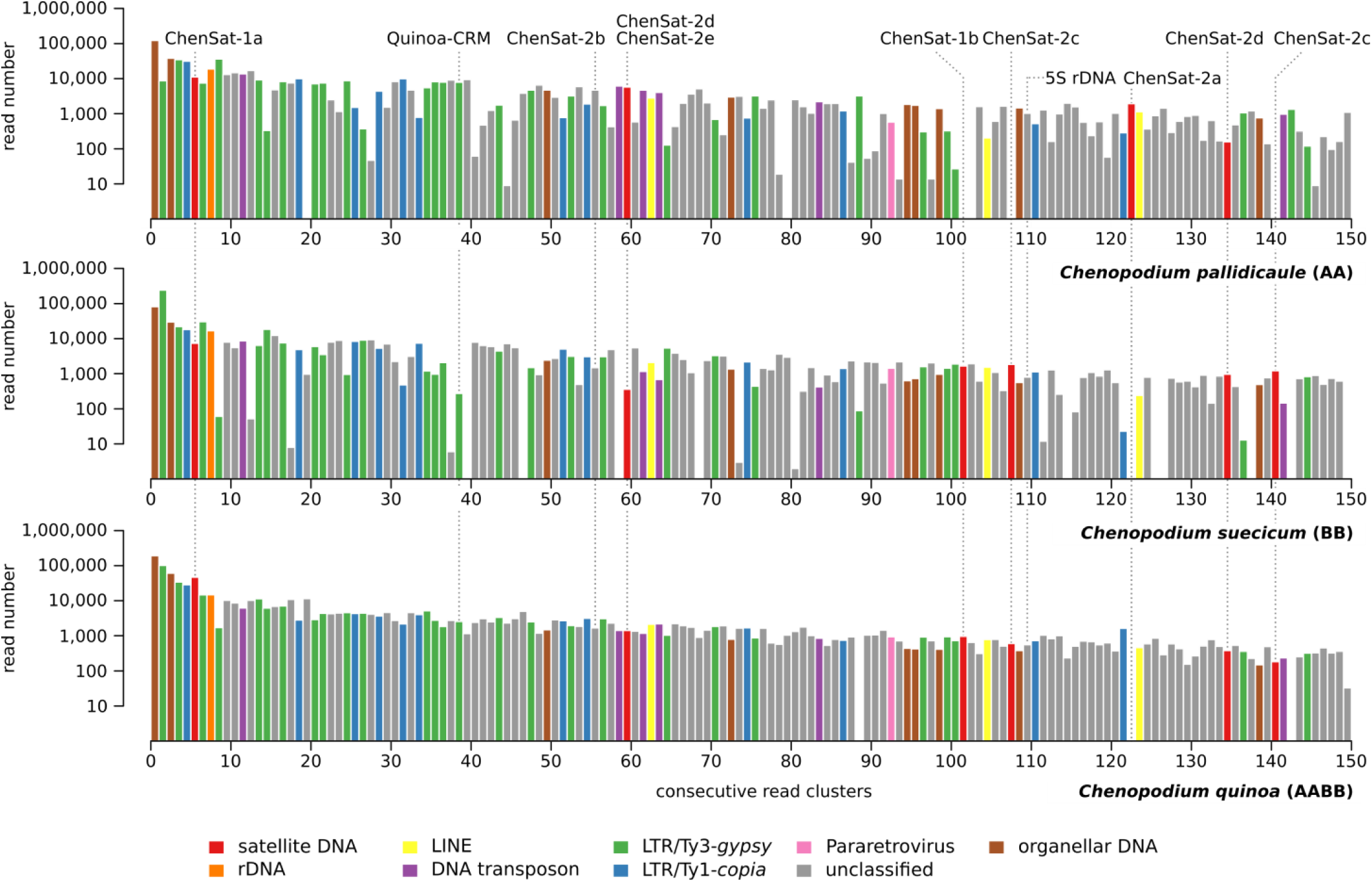
Comparative read cluster analysis of the most abundant repeats in the *C. pallidicaule*, *C. suecicum* and *C. quinoa* genomes. The barplots show the 150 most abundant *Chenopodium* read clusters sorted by decreasing genome contribution (X-axis). Identical cluster numbers refer to the same repeat families. Please note the logarithmic scale of the Y-axis. The read clusters were classified as indicated by the colour-code (see legend). Clusters corresponding to the analysed repeats are marked.

Based on star- or ring-like cluster shape and tandem arrangement within the cluster consensus sequence, we selected eight clusters harbouring seven *Chenopodium* satDNAs for further analysis. The sequences were grouped into families and subfamilies according to their sequence similarity, and designated ChenSat-1a, ChenSat-1b, ChenSat-2a, ChenSat-2b, ChenSat-2c, ChenSat-2d, and ChenSat-2e. Gaps in the bar plots indicate absence from a genome, such as seen for the satDNA families ChenSat-1b, ChenSat-2a, and ChenSat-2c (Figure 1). The number of reads indicates a satDNA fraction of 2.4 %, 1.5 % and 5.4 % for *C. pallidicaule*, *C. suecicum*, and *C. quinoa* relative to their overall repeat fractions. As read cluster abundance can misrepresent the real satDNA contribution (Novák et al., 2010, Ruiz-Ruano et al., 2016), we analysed the graphs to uncover clusters with masked satDNA (Figure S1, see explanation in Appendix S1). As a result, we directly inferred higher amplification in either A genomes (for ChenSat-2a, ChenSat-2b, and ChenSat-2e) or B genomes (for ChenSat-1b, ChenSat-2c, and ChenSat-2d).

### Assignment of *Chenopodium* satDNA to major families

To analyse the identified tandem repeats, we derived the monomer consensus sequences by independent and iterative mapping of sequence reads from each of the three genomes. This resulted in subgenome-specific reference sequences for satellite monomers and 5S rRNA genes including the spacer. The seven analysed satDNA consensus sequences from the three *Chenopodium* genomes fall into only two satDNA families, ChenSat-1 and ChenSat-2, marked by similar monomer lengths, conserved sequence stretches, and moderate pairwise sequence identities between 40 to 60 %.

For ChenSat-1a, the *C. quinoa*, *C. pallidicaule* and *C. suecicum* consensus monomers are highly similar with >95 % sequence identity. Its 40 bp monomers are marked by a very low G/C content (26-27 %) and make up a high genome proportion in all analysed genomes (0.6-

3.7 %, Table 1). ChenSat-1a is the major satellite of *Chenopodium* genomes (Figure 2A) and has already been cloned from the *C. quinoa* genome (Kolano et al., 2011, accession HM641822). Surprisingly, ChenSat-1a also has a 87 % similarity to the 40 bp satellite pBC1447 from the distantly related wild beet *Beta corolliflora* (Gao et al., 2000, accession AJ288880), an Amaranthaceae species from the Betoideae subfamily.

**Fig. 2:**
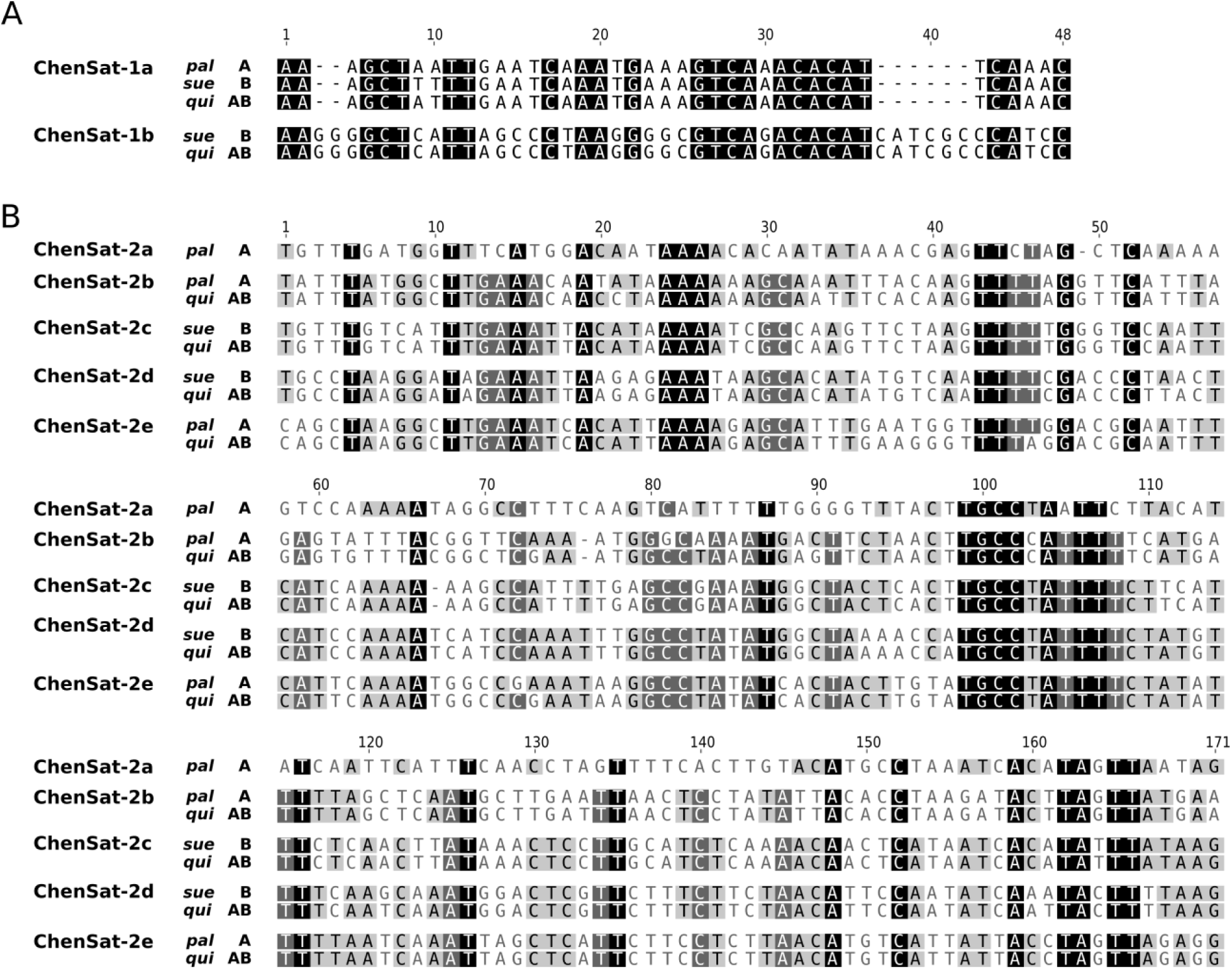
Species-specific consensus monomers of seven satDNAs from *C. pallidicaule* (*pal*), *C. suecicum* (*sue*), and *C. quinoa* (*qui*) genomes. **(A)** The alignment presents the genome-specific consensus sequences of ChenSat-1a and ChenSat-1b monomers, 40 and 48 bp in length, respectively. The genomes of *C. suecicum* (*sue*) and *C. quinoa* (*qui*) harbour both repeats, whereas *C. pallidicaule* (*pal*) contains ChenSat-1a, only. As visualised by the pairwise alignment, ChenSat-1a and ChenSat-1b share regions of high sequence identity (shaded in black) of 60 %. **(B)** Five variants of ChenSat-2 have been detected in the analysed *Chenopodium* species belonging to the same satellite superfamily. An alignment of species-specific consensus sequences from *C. pallidicaule* (*pal*, ChenSat-2a, -2b, -2e), *C. suecicum* (*sue*, ChenSat-2c, -2d), and *C. quinoa* (*qui*, ChenSat-2a to ChenSat-2e) genomes shows overall similar monomeric lengths of 170 to 171 bp, and regions of high sequence identity and similarity (shaded in black and grey).

**Table 1:** Tandem repeat characteristics

|  | Genome | Monomer length | GC content [%] | Pairwise identity [%] | No. of reads analysed † |
| --- | --- | --- | --- | --- | --- |
| ChenSat-1a | *C. pallidicaule* | 40 | 27 | 88 | 767 |
|  | *C. suecicum* | 40 | 27 | 86 | 2,479 |
|  | *C. quinoa* | 40 | 26 | 88 | 12,758 |
| ChenSat-1b | *C. suecicum* | 48 | 53 | 85 | 149 |
|  | *C. quinoa* | 48 | 53 | 84 | 387 |
| ChenSat-2a | *C. pallidicaule* | 170 | 31 | 93 | 2,338 |
| ChenSat-2b | *C. pallidicaule* | 170 | 30 | 87 | 781 |
|  | *C. quinoa* | 170 | 30 | 89 | 361 |
| ChenSat-2c | *C. suecicum* | 170 | 30 | 88 | 2,077 |
|  | *C. quinoa* | 170 | 30 | 86 | 679 |
| ChenSat-2d | *C. suecicum* | 171 | 32 | 90 | 931 |
|  | *C. quinoa* | 171 | 31 | 85 | 212 |
| ChenSat-2e | *C. pallidicaule* | 171 | 24 | 89 | 6,942 |
|  | *C. quinoa* | 171 | 29 | 84 | 848 |
| Chen5S | *C. pallidicaule* (AA) | 331 | 48 | 96 | 994 |
|  | *C. suecicum* (BB) | 318 | 43 | 95 | 1,041 |
|  | *C. quinoa* (A-derived) | 319 | 39 | 93 | 235 |
|  | *C. quinoa* (B-derived) | 317 | 35 | 99 | 348 |

† number of 2x1.7 million paired reads mapping to species-specific consensus

With 48 bp, ChenSat-1b monomers are slightly longer than ChenSat-1a (Figure 2A). From the satDNAs studied here, ChenSat-1b has the highest G/C content with 53 %, contributing many cytosine targets for potential DNA methylation (Table 1). It is only present in reads from B (sub)genome species, i.e. *C. suecicum* and *C. quinoa*, but absent from the A genome species *C. pallidicaule* (Figure 3). The short satDNA monomers of ChenSat-1a and ChenSat-1b contain stretches of sequence similarity and have an overall identity of 60 %, indicating an evolutionary relationship.

**Fig. 3:**
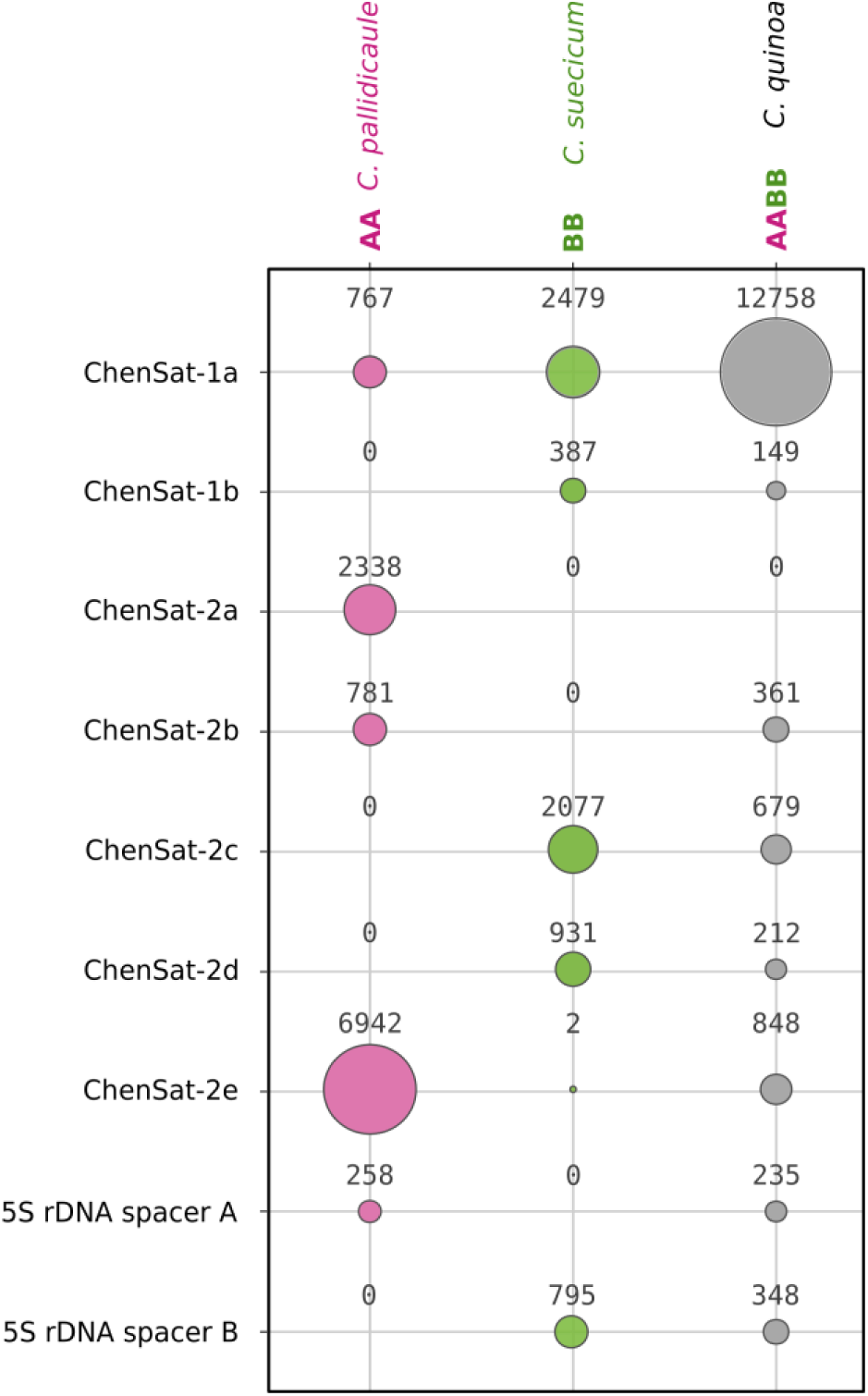
Quantification of tandem repeats within the *C. pallidicaule*, *C. suecicum* and *C. quinoa* genomes by read mapping. The bubble chart indicates relative abundance of the *Chenopodium* tandem repeats in the three analysed *Chenopodium* genomes. The filled area corresponds to the amount of reads mapping to each species-specific tandem repeat consensus. ChenSat-2a, ChenSat-2b, and ChenSat-2e are highly amplified in the A genome, with ChenSat-2a occurring solely in *C. pallidicaule* and not in the allotetraploid *C. quinoa* genome. Contrarily, ChenSat-2c, ChenSat-2d, and ChenSat-1b are amplified in the B genome with presence in *C. suecicum* and *C. quinoa* genomes. The 5S rDNA spacers also show a preference for A- or B-containing genomes, respectively.

Five of the *Chenopodium* satDNAs analysed here are diverged subfamilies forming the ChenSat-2 family. We hypothesize that ChenSat-2a to ChenSat-2e are derived from the same satDNA precursor with diversification during speciation. As visualised by a multiple sequence alignment and an all-against-all dotplot (Figure 2B, Figure S2), their monomer sequences share conserved residues over the whole length, with inter-family identities between 40.5 % and 59.6 %. ChenSat-2a to ChenSat-2e are highly amplified in only one of the two (sub)genomes, G/C contents below 32 %, and satDNA-typical monomer lengths of 170-171 bp (Table 1, Figure 3), presumably indicating selective constraints on the monomer size.

To test whether (sub)genome-specificity is restricted to non-coding tandem repeats, we derived four consensus 5S rDNA variants, one from *C. pallidicaule*, one from *C. suecicum*, and two from *C. quinoa*, derived from the A and B subgenomes, respectively. As expected, the 5S rRNA genes, including the regulatory boxes A, IE, and C, are 100 % identical in all analysed *Chenopodium* species (Figure S3), and differ only slightly from more distant plant 5S rRNA genes (e.g. 92 % identity to pine). However, between the subgenomes, the 5S rDNA spacers accumulated differences, including point mutations and a 12 bp insertion in *C. pallidicaule* (Figure S3). This sequence divergence is sufficient to assign the 5S rDNA spacers from *C. quinoa* to either the A or the B subgenome.

### Higher-order-structure, head-to-head junctions and retrotransposon association of quinoa satellites

Using 130,314 *C. quinoa* single molecule real-time (SMRT) reads, we searched for higher order arrangements and changes in repeat orientation. To achieve this, we identified tandem repeat-containing SMRT reads with a robust nHMM search (Table S1), retrieved 7,319 sequences and prepared dotplots for visual inspection. We observed four different patterns of tandem repeat organisation: continuous arrays, short arrays, inversions, and higher order arrangements. For all *C. quinoa* tandem repeats (ChenSat-1a, ChenSat-1b, and ChenSat-2b to ChenSat-2e, 5S rDNA), exemplary dotplots of 5,000 bp sequence stretches were shown, representative for each category (Figure S4). All analysed tandem repeats occur in both, long and short arrays. Head-to-head organisation was rare and only found in sequence reads containing ChenSat-1a, ChenSat-2b, and ChenSat-2e. Strikingly, most ChenSat-1a arrays identified were organised in higher order, with array lengths ranging from 1,000 bp to 2,000 bp, occasionally exceeding 5,000 bp. Similar HOR structures have been identified for ChenSat-1b, ChenSat-2c, and ChenSat-2e.

For ChenSat-1a, we observed dotplots with array interruptions. Inspection of the interrupting sequences revealed the interspersion with long terminal repeat (LTR) retrotransposons of different lineages. Most strikingly, ChenSat-1a arrays have been interrupted by Ty3-*gypsy* retrotransposons of the CRM clade (Figure S5). In order to assign this retroelement to the chromoviruses, we compared its key enzyme, the reverse transcriptase (RT), with other chromovirus RTs from Neumann *et al*. (2011). We clearly assigned this *Chenopodium* retrotransposon to the centromeric group A chromoviruses (Figure S6A), often marked by an integration preference for the centromeric heterochromatin (Neumann *et al*., 2011). It has a high similarity to the centromeric chromoviruses Beetle1, Beetle2 and Beetle7 in the related genera *Beta* and *Patellifolia* (Weber and Schmidt, 2009, Weber et al., 2013), all also known to co-localise with satDNA. An in-depth comparison of the integrase region enabled the identification of a C-terminal chromodomain with the CR-motif (Figure S6B), presumably conferring integration preference into centromeres (Novikova, 2009, Neumann et al., 2011). This retrotransposon is also highly repetitive and is represented by *RepeatExplorer* cluster CL39 (Figure 1).

### *Chenopodium* A and B genomes have distinct satDNA profiles

In order to quantify the tandem repeat abundance in the A and B genome diploid species, we individually aligned reads from each of the three genomes against the monomeric consensus sequences. Individual read mapping counts of *C. quinoa*, *C. suecicum*, and *C. pallidicaule* monomers (Figure 3) showed that ChenSat-1a is present in all three species, with a five to sixteen-fold copy number in *C. quinoa* as compared to the diploid genomes. ChenSat-2a has been exclusively detected in *C. pallidicaule*, and ChenSat-2b in *C. pallidicaule* and *C. quinoa*, suggesting their specificity for the A genome. Likewise, B-genome-specificity was inferred for ChenSat-1b, ChenSat-2c, and ChenSat-2d, all with hits in *C. suecicum* and *C. quinoa*, but absence in *C. pallidicaule*. ChenSat-2e has been detected in all three species, however with large read count differences, with a particular high amplification in the A genome diploid *C. pallidicaule* (6,942 reads) and the A chromosomes of *C. quinoa* (848 reads). The 5S rDNA is present in all genomes, however, the A- and B-genome-derived 5S rDNA spacers from quinoa, have been only detected in A and B genomes, respectively.

As next-generation sequencing is biased against G/C- and A/T-rich sequences, sometimes differing strongly from the genomic mean G/C values (Benjamini and Speed, 2012, Chen et al., 2013), quantification of satDNA based on read counts may misrepresent the genomic satDNA abundance. To verify the genome-specificity detected by bioinformatics (Figure 3), and to estimate the tandem repeat abundance in *Chenopodium* and related genera, we comparatively hybridised the ChenSat probes onto restricted genomic DNA (Figure 4). We tested eighteen species (Table 2), including two A genome diploids (lanes 1-2), two B genome diploids (lanes 3-4), two allopolyploids containing A and B subgenomes (lanes 5-6), distantly related *Chenopodium vulvaria* with the diploid V genome (lane 7), further related species from all sections of the Chenopodioideae (lanes 8-15), and three outgroups from the Betoideae (lanes 16-18). In order to release the satDNA-typical ladder-like patterns, restriction enzymes have been chosen for each tandem repeat as indicated in Figure 4.

**Fig. 4:**
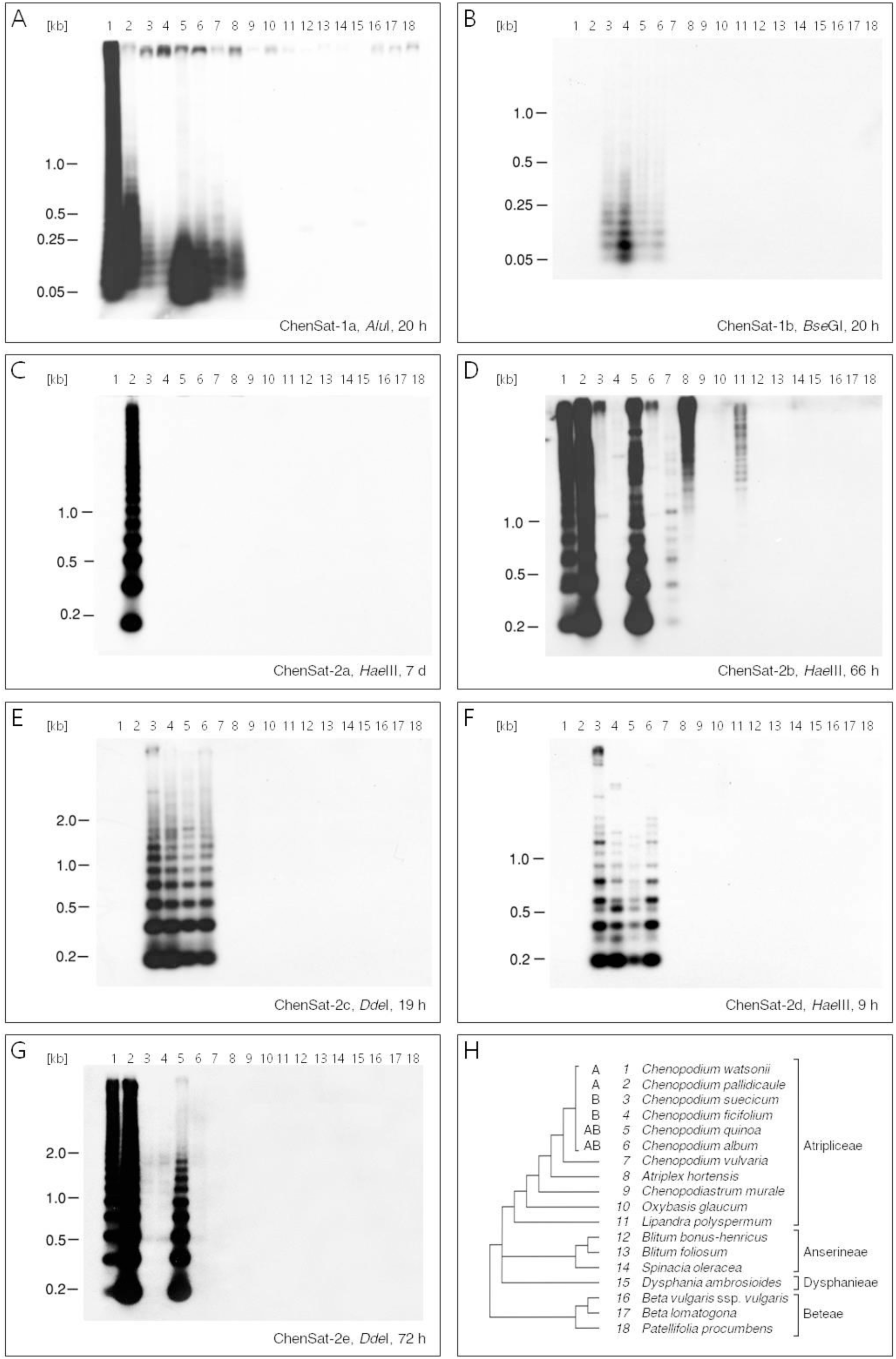
Abundance, specificity and genomic organisation of *Chenopodium* tandem repeats in the genus *Chenopodium* and in related plant genomes. Genomic DNA of eighteen plants has been restricted as indicated in each panel and was analysed by comparative Southern hybridisation of ChenSat-1a **(A)**, ChenSat-1b **(B)**, and ChenSat-2a to ChenSat-2e **(C-G)**. Exposure times ranged between nine hours and two weeks as indicated below the autoradiographs. **(H)** Selected plant species, their corresponding lanes, and their relationship according to published *trnL-F* and *matK/trnK* phylogenies (Fuentes-Bazan *et al*., 2012a). Branch lengths are not to scale.

**Table 2:**
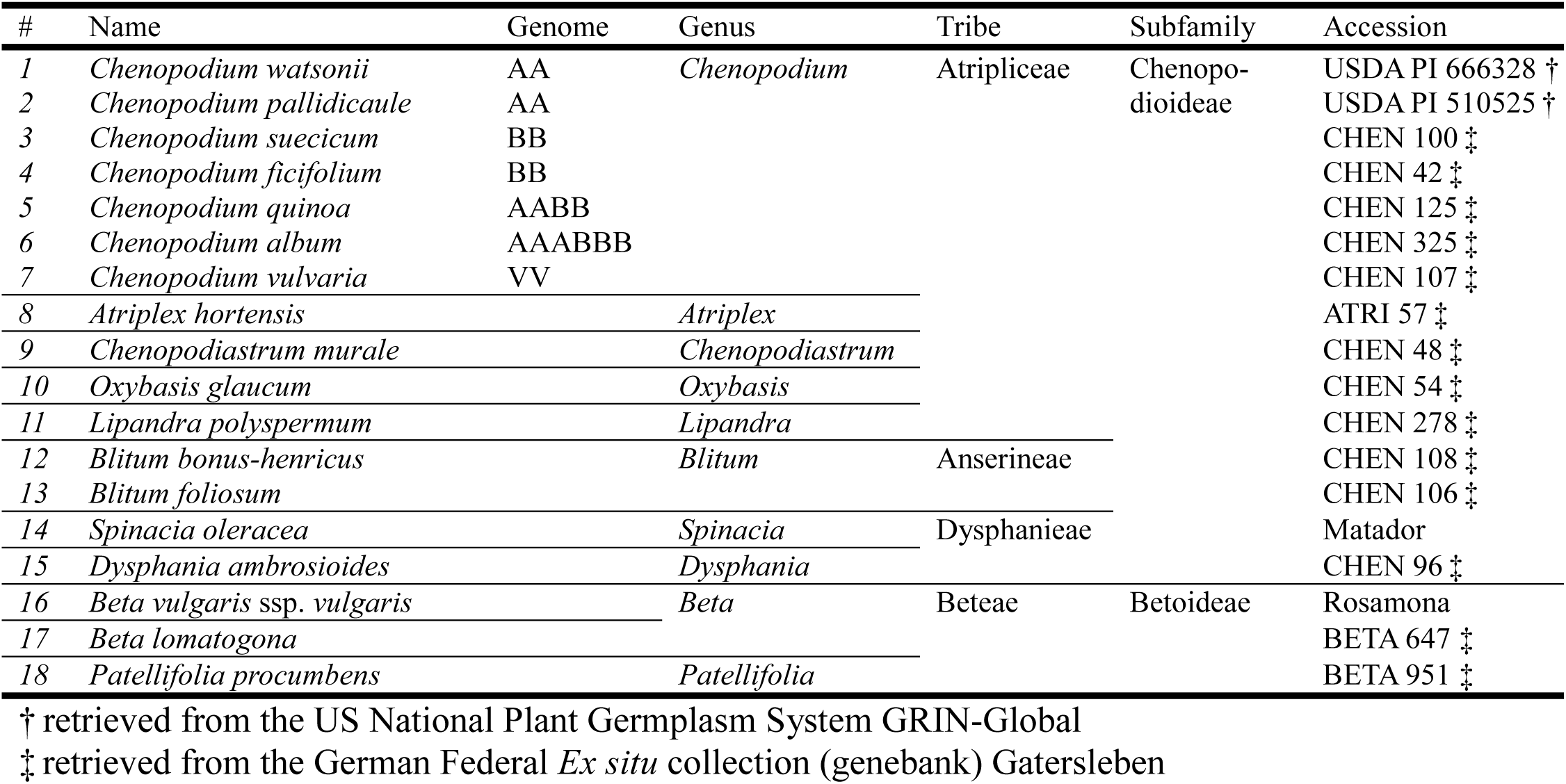
Plant material

ChenSat-1a occurs in all tested *Chenopodium* species (lanes 1-7) and in the closely related *Atriplex hortensis* (lane 8), producing ladder patterns with similar band sizes, consistent with the 40 bp monomer (Figure 4A). ChenSat-1a is highly abundant in A genome-containing species (*C. watsonii*, *C. pallidicaule*, *C. quinoa*, *C. album*) and abundant, but in lower copy numbers, in the B subgenome diploids (*C. suecicum*, *C. ficifolium*) and the more distantly related species (*C. vulvaria*, *A. hortensis*).

In contrast, ChenSat-1b shows only signals in the species containing B genomes, such as *C. suecicum*, *C. ficifolium*, *C. quinoa* and *C. album* (Figure 4B, lanes 3-6). The hybridisation generates similar restriction ladders for all four species, with the highest abundance in the diploid *C. ficifolium* (lane 4). The autoradiogram verifies the short ChenSat-1b monomer length (48 bp), with bands up to the hexamer and a smear ranging up to 500 bp.

ChenSat-2a hybridises exclusively to *C. pallidicaule* (Figure 4C, lane 2), and is absent from the other A genome containing di- and polyploids tested. This is in line with its exclusion from the *C. quinoa* and *C. suecicum* genomes as detected by read mapping and clustering (Figure 3, Figure S1). Hybridisation yields ladders with strong monomeric bands, supporting the 170 bp monomer length, and signals up to the decamer, before falling together to form a smear.

We detected a strong ladder-like hybridisation pattern of the ChenSat-2b subfamily for both A genome diploids (Figure 4D, lane 1, 2) and *C. quinoa* (lane 5). The *C. vulvaria* genome produces weak ladder hybridisation patterns, indicating presence of ChenSat-2b (as the only repeat from the ChenSat-2 family) in more distantly related *Chenopodium* species. The banding pattern supports a conserved ChenSat-2b monomer length of 170 bp (lanes 1, 2, 5, 7). Interestingly, *C. suecicum* (lane 3), *C. album* (lane 6), and *Lipandra polyspermum* (lane 11) produce moderate, and in particular *A. hortensis* (lane 8) produces strong signals in the high molecular weight fraction of the DNA. This indicates presence of diverged repeats lacking the conserved *Hae*III site in these genomes, presumably also belonging to ChenSat-2b or a closely related subfamily.

Similar to ChenSat-1b, the B-specific ChenSat-2c and ChenSat-2d probes hybridise exclusively to the B-genome-containing species *C. suecicum*, *C. ficifolium*, *C. quinoa*, and *C. album* (Figure 4B, E, F, lanes 3-6). ChenSat-2c hybridisation generates ladder patterns in all four species, without monomer length variation (Figure 4E). We observe equally strong and similarly spaced ladder patterns up to the nonamer and higher. Hybridisation with ChenSat-2d produces two superimposed restriction ladders for all B-subgenome-containing species, indicating presence of an internal *Hae*III site in some monomers (Figure 4F). Indeed, apart from a single canonical *Hae*III site, the ChenSat-2d consensus sequence contains five additional positions, in which a single nucleotide mutation could result in an intact 5’-GGCC-3’ restriction site (Figure 2). *C. quinoa* (lane 5) shows a weaker hybridisation pattern, likely caused by reduced ChenSat-2d abundance and consistent with the read mapping.

We detected strong ladder-like hybridisation of ChenSat-2e to the A genome diploids (lane 1-2) and to *C. quinoa* (lane 5) after short exposure (not shown). After long exposure (72 hours), weaker signals with reduced ladder patterns in B diploids were detectable (Figure 4G), indicating very low abundance in B-containing genomes.

Summarising, for the allotetraploid *C. quinoa*, and the diploids *C. pallidicaule* and *C. suecicum*, we collected computational and experimental evidence for differential satDNA amplification in A and B genomes, and consistently provide an in-depth view into the satDNA distribution in Chenopodioideae genomes.

### Satellite DNA allows the assignment of chromosomes to the A and B subgenome

To determine the satDNA localisation along chromosomes we hybridised the 5S and the 18S-5.8S-25S rRNA genes as well as the newly identified tandem repeats to *C. pallidicaule* (AA diploid), *C. ficifolium* (BB diploid) and *C. quinoa* (AABB tetraploid) metaphase spreads (Figure 5). For assignment of the *C. quinoa* chromosomes to either the A or the B subgenome, we re-hybridised five metaphases with genomic DNA of B-derived *C. suecicum* by GISH (Figure 5 E,I,K,M,O, green).

**Fig. 5:**
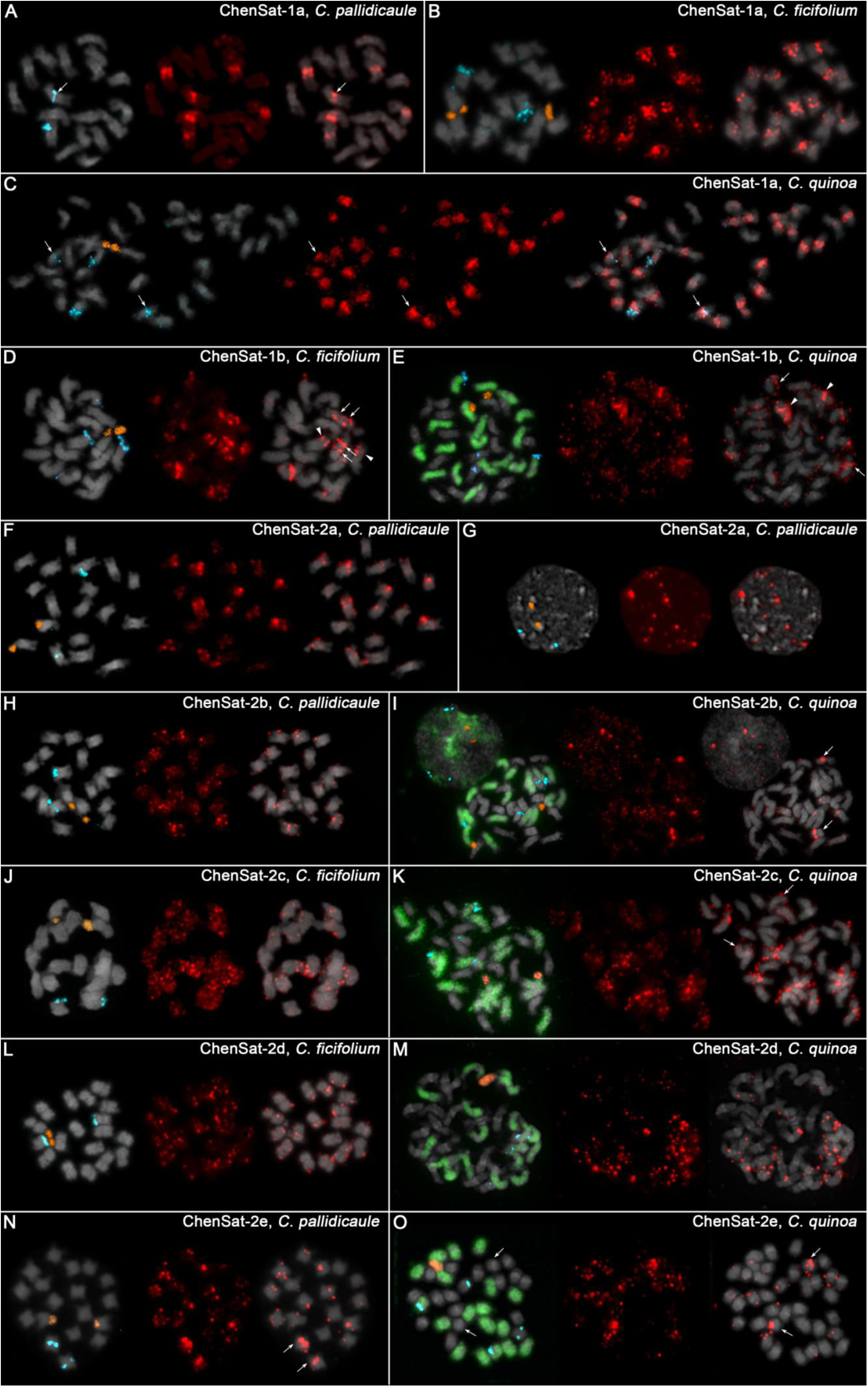
Multi-colour fluorescent and genome *in situ* hybridisation to chromosome spreads of *C. quinoa*, *C. pallidicaule*, and *C. ficifolium*. DAPI-stained mitotic chromosomes are shown in grey. We coupled biotin to all satellite DNA probes (red signals), specifically ChenSat-1a **(A-C)**, ChenSat-1b **(D-E)**, ChenSat-2a **(F-G)**, ChenSat-2b **(H-I)**, ChenSat-2c **(J-K)**, ChenSat-2d **(L-M)**, and ChenSat-2e (**N-O**). The 18S-5.8S-25S rDNA (orange) and 5S rDNA (light blue) were co-hybridised for easier chromosome identification. For the allotetraploid *C. quinoa* spreads **(panels E, I, K, M, O)**, we additionally hybridised *C. suecicum* genomic DNA (green signals) to allocate B-genome-derived chromosomes. The arrows and arrowheads are specified in the man text.

Regarding the 5S rRNA genes, in *C. pallidicaule*, we observed a major signal on two chromosomes in the interstitial region (Figure 5A,F,H,N, blue). This is in line with previous reports for the 5S rDNA of *C. pallidicaule* (Kolano *et al*., 2012). In *C. ficifolium*, hybridisation of the 5S rRNA genes results in a distal signal on one chromosome pair (Figure 5B,D,J,L, blue). In tetraploid *C. quinoa*, we observed strong 5S rDNA signals on four chromosomes (Figure 5C,E,I,K,M,O, blue). Whereas the two B-derived chromosomes carry a distal signal, the two A-derived chromosomes harbour the 5S rDNA in the interstitial region (Figure 5E,I,O), also in line with prior reports (Maughan et al., 2006, Kolano et al., 2012). Hybridisation of the 18S-5.8S-25S rRNA genes revealed a single chromosome pair with a distal signal on one chromosome arm for all three genotypes, despite *C. quinoa*’s tetraploidy (Figure 5, panels B-F, H-O, orange). GISH to *C. quinoa* metaphases indicated the origin of the 18S-5.8S-25S rRNA genes from B genomes (Figure 5E,I,K,M,O), corroborating previous reports (Maughan et al., 2006, Kolano et al., 2016).

As ChenSat-1a has been detected in all three genotypes as visible by Southern hybridisation (Figure 4A), it was hybridised to chromosome spreads of each of the three accessions (Figure 5A-C). In the *C. pallidicaule* A genomes, six strong, pericentromeric ChenSat-1a signals (Figure 5A, red) were detected on metaphase spreads, including the chromosomes harbouring the 5S rRNA genes (Figure 5A, blue signals). ChenSat-1a is positioned in close vicinity and partially overlapping to the interstitial 5S rRNA gene array on one chromosome (Figure 5A, arrowed). In the B genome of *C. ficifolium* ChenSat-1a produces eight strong signals, often localising in the pericentromeric regions and spreading into intercalary regions of the remaining chromosomes (Figure 5B, red). Co-localisation of ChenSat-1a with 5S and 18S-5.8S-25S rRNA genes has not been observed (Figure 5B, blue, orange).

ChenSat-1a was detected in the pericentromeric chromatin of all *C. quinoa* chromosomes with 24 major, 8 moderate and 4 minor signals (Figure 5C, red). We observed co-hybridisation of ChenSat-1a and two of the 5S rRNA gene signals (Figure 5C blue, arrows), verified by presence of both repeats on identical SMRT reads (Table S2). The chromosome pair carrying the distal 18S-5.8S-25S rDNA contained pericentromeric, moderate ChenSat-1a signals.

In *C. ficifolium*, the chromosome pair with the 18S-5.8S-25S rRNA genes (orange) carries two major ChenSat-1b signals in the intercalary and pericentromeric regions (Figure 5D, arrows). The distal 5S rRNA genes (blue) mark an additional chromosome pair with major, intercalary ChenSat-1b signals on one chromosome arm (Figure 5D, red, arrowheads). We observed further major signals in intercalary and pericentromeric regions of a third chromosome pair (Figure 5D). Similarly, ChenSat-1b is localized along distal and pericentromeric regions of the 18 chromosomes of the *C. quinoa* B subgenome (Figure 5E, green), with two major and four moderate signals, indicating large ChenSat-1b arrays (Figure 5E). Similar to *C. ficifolium* (Figure 5D), the two major signals are located on the 18-5.8-28S rDNA-carrying chromosome pair, whereas two of the four moderate signals can be found in the intercalary regions of the chromosome arms carrying the distal 5S rDNA (Figure 5E, arrowheads, orange and arrows, blue signals).

ChenSat-2a is present on all *C. pallidicaule* chromosomes with varying intensities (Figure 5F). It is localised mainly in the interstitial, but also in the distal chromosomal areas. At the higher resolution of interphase nuclei, we observed that ChenSat-2a is largely excluded from the strongly DAPI-positive heterochromatin (Figure 5G).

In *C. pallidicaule*, the ChenSat-2b signals are dispersed on all chromosomes with varying signal intensities, mostly in interstitial and intercalary regions (Figure 5H). In *C. quinoa*, ChenSat-2b is dispersed on many of the 18 A-genome-derived chromosomes (Figure 5I, unlabeled chromosomes). Two chromosomes, presumably a pair, carry a major intercalary signal on one arm (arrows), suggesting presence of large ChenSat-2b arrays.

ChenSat-2c produces weak to moderate signals and is distributed in the intercalary and distal regions of the *C. ficifolium* chromosomes (Figure 5J, red). Only two chromosomes show stronger hybridisation. As indicated by FISH and GISH on *C. quinoa* metaphases, ChenSat-2c is present on all 18 B-subgenome-derived chromosomes (Figure 5K). These signals are present in low to moderate signal strength in intercalary and pericentromeric regions. We observed ChenSat-2c co-localisation with the distal and B-genome-derived 5S rDNA (arrows).

On chromosomes of *C. ficifolium*, ChenSat-2d is widely dispersed on chromosomes with low to moderate intensities (Figure 5L). In *C. quinoa* only B-genome-derived chromosomes (Figure 5M green) showed ChenSat-2d arrays of different size distributed in the intercalary and distal chromosome regions, with exclusion from the pericentromeric regions.

At *C. pallidicaule* metaphases, ChenSat-2e produces two very strong hybridisation sites on the 5S rDNA-carrying chromosome pair (Figure 5N, blue, arrowed). Twelve chromosomes show weak to moderate signals and four chromosomes give only faint signals. Similarly, in *C. quinoa*, a chromosome pair carries a major signal (Figure 5O, arrows), however, without co-localisation to the 5S rRNA genes. The GISH reveals that the two major signals localise on A-genome-derived chromosomes (arrows).

### Intermingling of tandem repeat families along *C. quinoa* chromosomes as detected on SMRT reads, scaffolds and by FISH

To infer evolutionary relationships and potential exchange between homoeologous chromosomes, we analysed the physical neighbourhood and the intermingling of tandem repeats on *C. quinoa* SMRT reads, scaffolds, and (pseudo)chromosomes. Using a tandem repeat nHMM analysis, we found that 201 out of 130,314 *C. quinoa* SMRT reads harboured arrays from at least two different satellite repeats (Table S2). We focused on two combinations:

I. ChenSat-1a and ChenSat-1b were detected on six reads (Figure 6A). As both repeats share considerable sequence identity (Figure 2A) and are potentially related, we verify their co-localisation by multi-colour FISH with ChenSat-1a and ChenSat-1b probes. We identified arrays of both tandem repeats in the pericentromeric region of two metaphase chromosomes (Figure 6C arrows) and showed interspersion along stretched chromatin fibres (Figure 6D).
II. Combinations of ChenSat-2c and ChenSat-2e are detected on 91 SMRT reads (Figure 6B) with both satellites organised in short arrays. ChenSat-2e is strongly enriched in the A genome, whereas ChenSat-2c is restricted to B-derived regions. Yet, using dual-colour FISH, we identified both repeats closely associated on the same chromosome pair (Figure 6E, arrows).

**Fig. 6:**
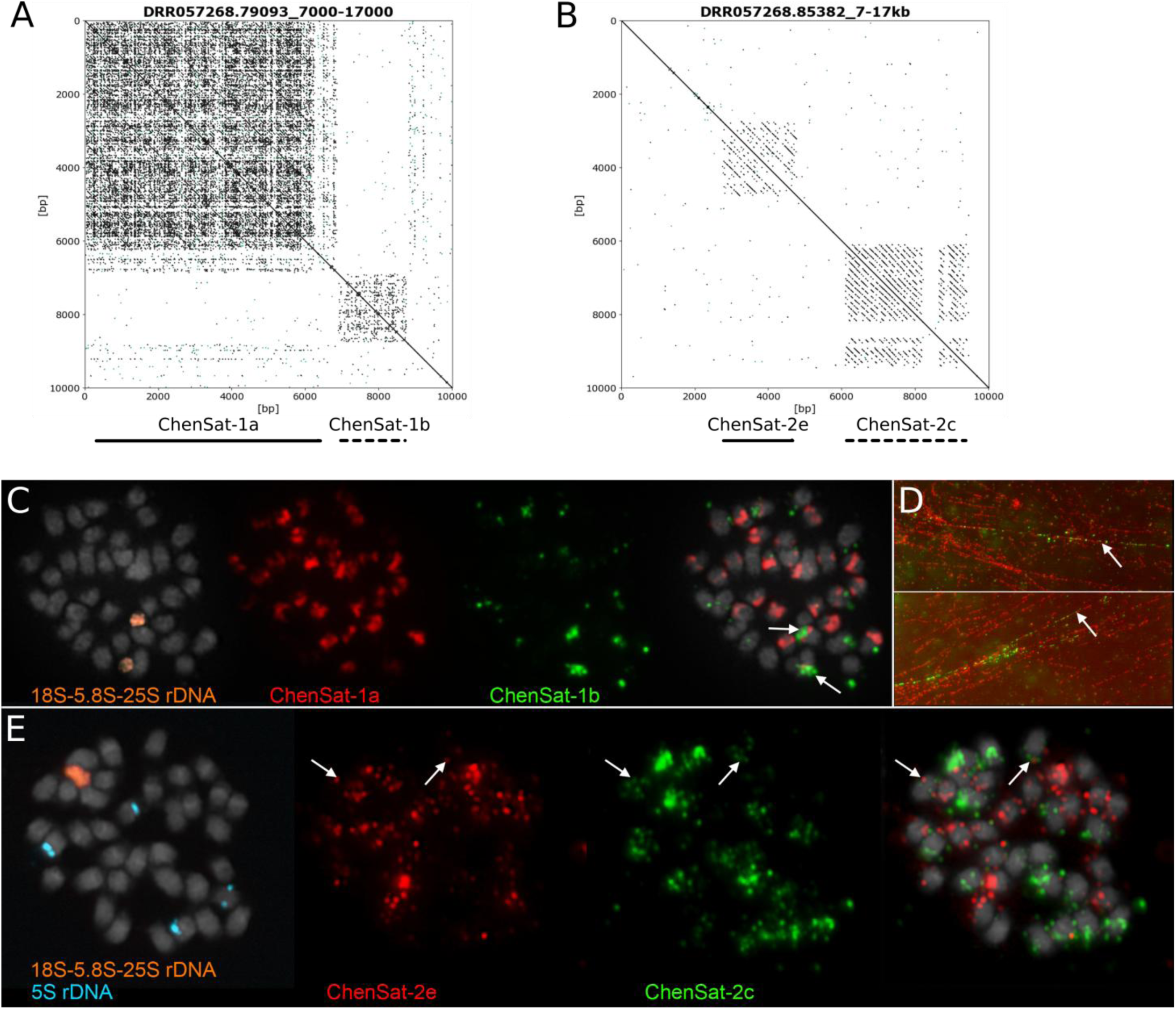
Co-localisation of satDNAs in the *C. quinoa* genome investigated on long reads and by cytogenetics. **(A)** ChenSat-1a and ChenSat-1b arrays have been detected on six SMRTs reads. Here, a dotplot of a representative 10 kb region from a SMRT read is shown. **(B)** Similarly, short arrays of ChenSat-2c and ChenSat-2e have been detected next to each other on SMRT reads in 91 cases. A dotplot of a 10 kb region of a representative read is shown. **(C)** Dual-colour FISH on *C. quinoa* metaphase chromosomes provides practical evidence that ChenSat-1a (red) and ChenSat-1b (green) co-occur on two chromosomes (arrows). We showed DAPI-stained chromosomes and fibres in grey, and additionally probed the 18S-5.8S-26S rDNA (orange) for easier chromosome allocation. **(D)** Hybridisation of ChenSat-1a (red) and ChenSat-1b (green) to stretched fibres further supports their interspersed arrangement. **(E)** Dual colour-FISH of ChenSat-2c (red) and ChenSat-2e (green) on *C. quinoa* metaphases.

As the 5S rRNA gene variants from the *C. quinoa* A and B genomes differ strongly in their spacer sequences (Figure 7A), they can also be used to detect interlocus homogenisation. However, the nHMM approach did not detect any co-occurrence of A- and B-derived 5S rRNA genes. To verify this, we exemplarily selected the eight longest *C. quinoa* SMRT reads, which were completely covered by 5S rDNA tandem repeats, and extracted 283 genes with spacer. The 5S rDNA monomers were aligned and their relationship was visualised by a dendrogram (Figure 7B). All monomers fall into one of two groups, belonging to either the A or B genome. All A-genome-specific monomers have been derived from three of eight reads, whereas all B-genome-specific monomers have been extracted exclusively from the remaining five SMRT reads (see magenta and green line colour in Figure 7B, respectively). This indicates maintenance of the 5S rRNA gene arrays without exchange between homoeologous chromosomes of the quinoa A and B genome.

**Fig. 7:**
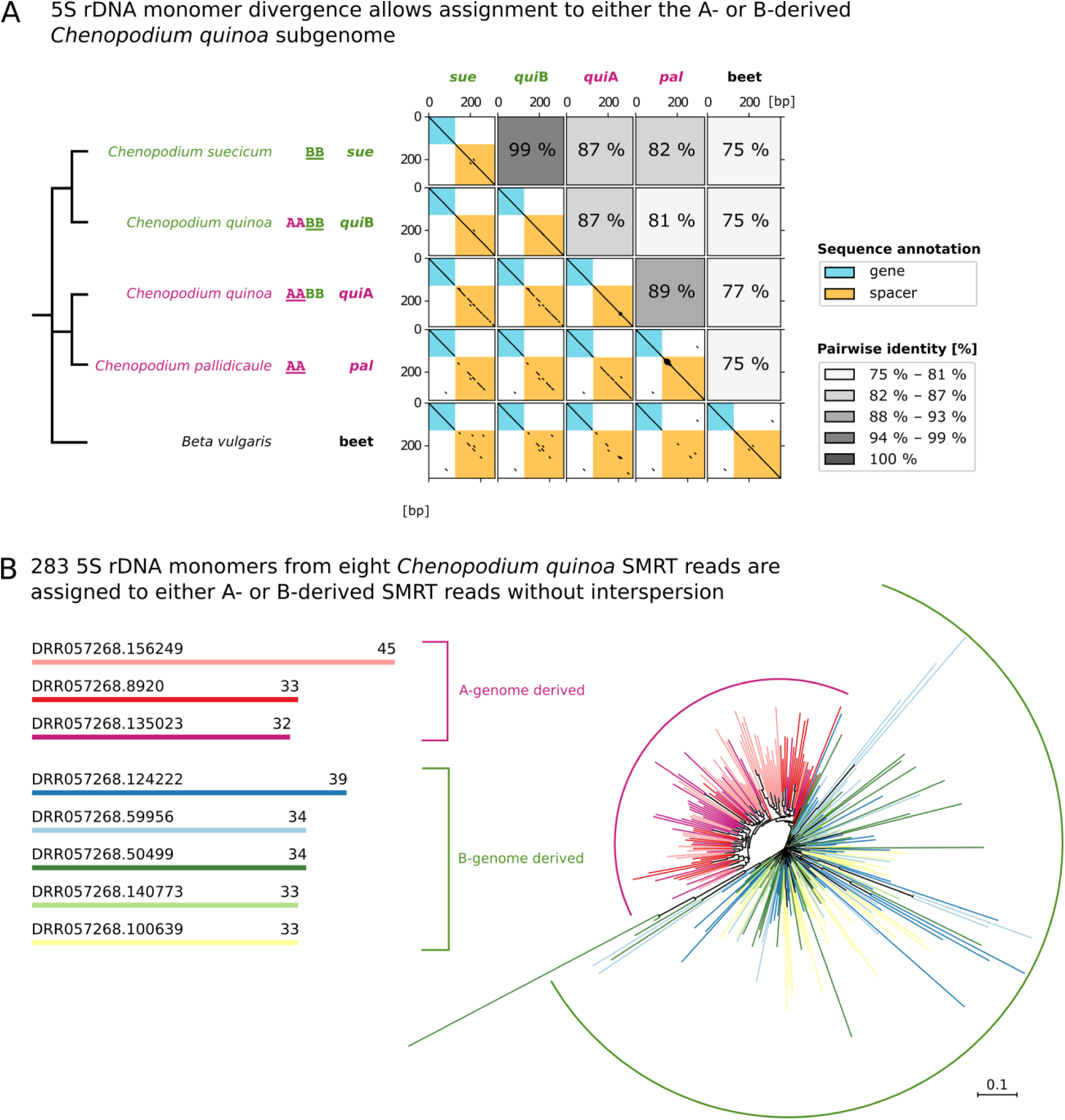
The analysed *Chenopodium* genomes contain homogeneous 5S rDNA arrays with either A or B genome variants. **(A)** Divergence of the spacer from the 5S rDNA allows the monomer assignment to the A or B subgenome from *C. quinoa*. Dotplot visualisation of the four 5S rDNA consensuses shows high similarity among the A-derived (*pal* and *qui*A) as well as among B-derived (*sue* and *qui*B) 5S rDNA monomers. The conserved gene (turquoise) and the variable spacer region (ochre) are indicated by shading. A comparison between (sub-)genomes, shows the accumulation of single nucleotide polymorphisms and indels, resulting in lower identities, which range between 81 % and 87 %. The dotplot was generated using a wordsize of 10 with toleration of 1 mismatch. For the corresponding multiple sequence alignment see Fig. S3. **(B)** Homogeneity of 5S rDNA arrays was analysed by extraction of 283 5S rDNA monomers from eight *C. quinoa* SMRT reads. We considered each SMRT read as an individual array. The monomers have been aligned and grouped by a Neighbor-Joining analysis. As 5S rDNA monomers derived from A and B subgenomes differ vastly in their spacer regions, the resulting dendrogram forms two major branches, each representative for A- and B-derived variants. For three reads, all 110 monomers group exclusively with the A subgenome reference, whereas the remaining five reads (173 monomers) were assigned to the B subgenome group. We did not find evidence for intermingling of 5S rDNA variants, and no indication of interlocus recombination between homoeologous chromosomes.

Although some SMRT reads can span up to 50 kb, they do not scale to a chromosome. For a larger overview, we used 619 high-quality scaffolds from the *C. quinoa* genome sequence (Jarvis *et al*., 2017) to provide evidence of the interspersion and intermingling of tandem repeats. Based on gene data, 226 of the 619 scaffolds could now be assigned to either the A or B genome (Jarvis et al., 2017). If our genome-specific satDNAs were used as markers, 51 previously unassigned quinoa scaffolds have now been classified as either belonging to the A (24) or B (27) genome (Table S3). In six cases, we mapped both A- and B-genome-derived satDNAs to the same scaffold, indicating exchange between quinoa A and B genome. However, as these six cases could not be verified on the shorter SMRT reads, we cannot rule out incorrect genome assembly.

Taking together, we detected intermingling of several satDNA families along the chromosomes, using long reads, detection on scaffolds, and FISH. We co-localised both ChenSat-1 families in *C. quinoa*, which may be an indication of their common decent. For the ChenSat-2 subfamilies, sequence data may indicate DNA exchange between A and B subgenomes, whereas we can confidently exclude dispersal of the 5S rRNA genes in the respective other subgenome.

## DISCUSSION

### Next- and third-generation sequence reads give an overview on satDNA landscapes in *Chenopodium*

By clustering of next-generation sequence reads, we detected and classified the repetitive genome proportion of allotetraploid *C. quinoa* and the two potential progenitor genomes *C. pallidicaule* and *C. suecicum*, revealing repeat fractions of 73.2 %, 76.9 %, and 78.1 %, respectively. We consider these values as conservative estimates, and likely underestimations, as they do not account for the high rate of repeated DNA divergence. Repeats may have escaped their detection by accumulation of mutations over time, recombination, diversification, reshuffling, and decay (Ma et al., 2004, Wollrab et al., 2012, Elliott and Gregory, 2015, Sanchez et al., 2017, Bourque et al., 2018). From the extracted repeat dataset, we identified and characterised seven *Chenopodium* satDNAs and the 5S rRNA genes. Six out of the seven ChenSat repeats have been specific or enriched in either the *Chenopodium* A (ChenSat-2a, ChenSat-2b, ChenSat-2e) or B genome (ChenSat-1b, ChenSat-2c, ChenSat-2d), respectively. Only ChenSat-1a has been detected in both A and B genomes in high abundance.

Read cluster analyses of high-throughput data have already been used to gain access to tandem repeat families in a number of plants, such as pea, bean, onion, camellia, crocus, pepper, and fern (Macas *et al*., 2007, Heitkam *et al*., 2015, Kirov *et al*., 2017, Ávila Robledillo *et al*., 2018, Schmidt *et al*., 2019, Zhou *et al*., 2019), with genomic satDNA fractions ranging from 0.1 % to 36 % (Garrido-Ramos, 2017). In the allopolyploid *C. quinoa*, satDNA accounts for 5.4 % of the repeated DNA fraction and for 4.0 % of the genome. Compared to the closely related sugar beet with a satDNA proportion of 11.15 % (Kowar et al., 2016), the satDNA content of quinoa is low. However, close taxonomic relationship between species is not correlated with similarly sized repeat fractions; even within species of a single genus, satDNA families can amplify with vast differences as observed between *Fritillaria* species (Kelly et al., 2015).

Genome assemblies are error-prone, in particular in repetitive regions. For example, the human reference genome is one of best-studied genome assemblies, but still contains many inaccuracies regarding satDNA array length and higher order organisation (Miga, 2015, Jain et al., 2018). For non-model organisms, assemblies often contain even less information making the study of repetitive regions laborious (Peona et al., 2018). The availability of third-generation long reads opens the way to solve genomic and evolutionary questions targeting satellite and ribosomal DNAs, such as array length, abundance and organisation in higher-order structures or head-to-head arrangements (Sevim *et al*., 2016, Khost *et al*., 2017, Symonová *et al*., 2017, Lower *et al*., 2018, Cechova *et al*., 2019, Vondrak *et al*., 2019). For *C. quinoa*, we identified arrays of the satellite families ChenSat-1a, ChenSat-1b, ChenSat-2b, ChenSat-2c, ChenSat-2d, and ChenSat-2e on SMRT reads; all were arranged in short or long homogeneous arrays, but also in structures of higher order (ChenSat-1a, ChenSat-1b, ChenSat-2c, and ChenSat-2e) and head-to-head arrangements (ChenSat-1a, ChenSat-2b, and ChenSat-2e). Similar to observations in the pike genome (Symonová et al., 2017), we also detect inversions in the 5S rDNA, and long as well as short arrays. Higher-order arrangement of ChenSat-1a has also been detected on clones of multimers as reported very recently (Belyayev et al., 2019).

### Seven *Chenopodium* satDNAs fall into only two major satDNA families

We identified seven *Chenopodium* satDNAs, belonging to one of two families, the ChenSat-1 family with short 40-48 bp monomers or the ChenSat-2 family with 170-171 bp monomers.

The most abundant satDNA family in the analysed *Chenopodium* genomes is ChenSat-1a, initially published as *C. quinoa* clone 12-13j (Kolano et al., 2011, Orzechowska et al., 2018). Its short monomer size of 40 bp is unusual for a tandem repeats of high abundances, which usually consist of 160-180 bp or 320 to 360 bp monomers (Hemleben et al., 2007). SatDNAs have been split into micro- (2-5 bp monomer), mini- (6-100 bp monomer), and conventional satellites (>100 bp monomer) (Vergnaud and Denoeud, 2000, Mehrotra and Goyal, 2014). In recent times, the focus has shifted to recognise not only the satDNA’s monomer size, but its genomic organisation of repeating units into long arrays as the main molecular hallmark of satDNA (Richard et al., 2008). These long arrays have been verified for ChenSat-1a and ChenSat-1b, and we therefore do not consider them as minisatellites, but as canonical satellite DNA. Major satDNA arrays, made up of short monomers < 100 bp, have been occasionally observed in other plants: In *Ricinus communalis* a satellite with 39 bp monomers constitutes 8.2 % of the genome (Melters et al., 2013). Likewise, a 43 bp satellite from *Camellia japonica* occupies more than 11 % of the genome and is located distally on all chromosomes (Heitkam et al., 2015). An extreme case is the fern *Vandenboschia speciosa*, in which the numerous satDNAs have short monomer lengths (between 33 and 141 bp, Ruiz-Ruano et al., 2018). This underpins that satellites with short monomer lengths can form very large arrays as observed here for ChenSat-1a and ChenSat-1b.

Despite the short monomer length, we observed a ChenSat-1a organisation with and without higher order, indicating an evolution towards longer and more complex repeated ChenSat-1a patterns (as reviewed by Plohl *et al*., 2012). The ChenSat-1a higher order repetitions have not been of conserved lengths, but differ in size on the different long reads. This can explained by the hypothesis of Rudd et al. (2006), that higher-order and monomeric satDNAs evolve at different rates, leading to less conserved higher-order repeat units compared to monomers.

ChenSat-1a localises at the centromeric constriction of all *C. quinoa* centromeres. This is in line with the observation, that in most organisms, the main satellite is expected at the centromere (Jiang et al., 2003, Melters et al., 2013). Its arrays are interspersed with full-length chromoviruses of the CRM lineage, as described for many plant centromeres (Cheng et al., 2002, Weber et al., 2013). This chromovirus family was different from a previously characterised partial *C. quinoa* chromovirus reverse transcriptase of the Tekay clade (Kolano et al., 2013). Instead, it represents a canonical chromovirus of the A group, containing a chromodomain of the CR-type, typical for centromeric retrotransposons (Novikova, 2009, Weber and Schmidt, 2009, Neumann et al., 2011), thus providing support for ChenSat-1a’s association with centromeres.

Similar to ChenSat-1a, the related family ChenSat-1b has also a short monomer length of 48 bp. High sequence identities of 60 % over the whole length indicate a common decent. As ChenSat-1b has a different species distribution, limited solely to B-genome-derived *Chenopodium* genomes, we assume that ChenSat-1a is more ancient and likely the progenitor. Only in a few instances, we localized ChenSat-1b to the *C. quinoa* pericentromeric regions, making a role in the formation of active centromeres unlikely. This contrasts with observations for some plants, such as the common bean, in which genomes active centromeres are formed by two alternative satDNAs, CentPv1 or CentPv2 (Iwata *et al*., 2013).

The satDNA subfamilies ChenSat-2a, ChenSat-2b, ChenSat-2c, ChenSat-2d, and ChenSat-2e are characterised by the conventional monomer lengths of 170 and 171 bp, containing conserved sequence stretches, and pairwise sequence identities of about 60 %. Therefore, we consider them as subfamilies of the ChenSat-2 family, and postulate a common origin. All ChenSat-2 satDNA families form long arrays, as verified by Southern hybridisation and SMRT read analysis.

### Different repeat landscapes emerged during *Chenopodium* speciation into species with A and B genomes

Repeats are subject to rapid changes during adaptation to new environments, contributing to rapid genome evolution and speciation (Oliver et al., 2013, Stapley et al., 2015, Serrato-Capuchina and Matute, 2018). After allopolyploidisation, it is assumed that an imbalance of repeats and their epigenetic impact drive the polyploidisation life circle and often lead to genome size shrinkage, diploidisation, and subgenome dominance (Edger et al., 2017, Vicient and Casacuberta, 2017, Mhiri et al., 2019). We traced the satellite DNA evolution in the genus *Chenopodium* and follow the evolutionary history of *C. quinoa*, as summarised in our evolutionary scenario (Figure 9).

**Fig. 8:**
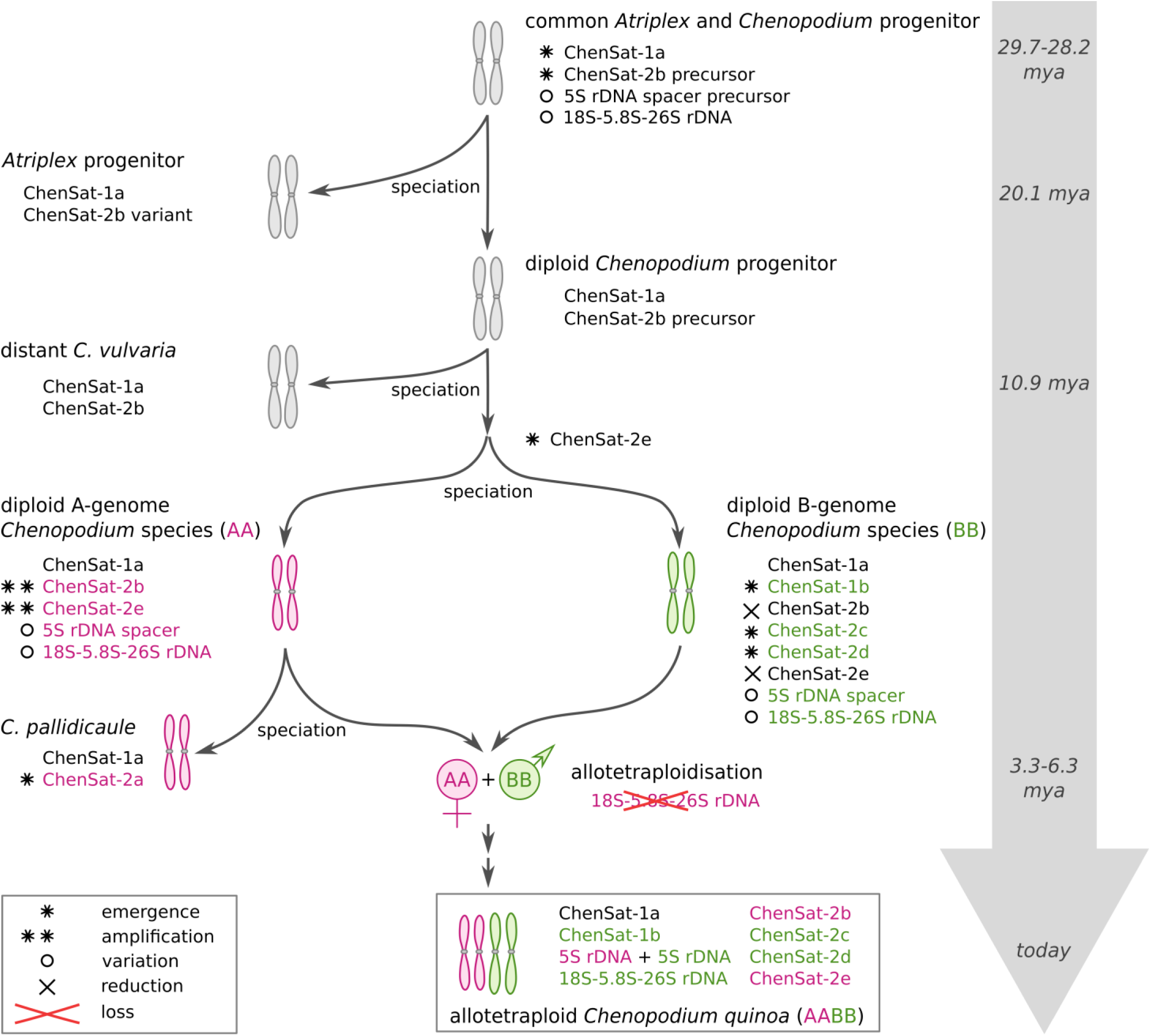
Scenario for the evolution of tandemly repeated DNA during the history of *Chenopodium* speciation and allopolyploidisation. The time axis includes estimates of the stem age of the Atripliceae (Kadereit et al., 2010), the early splits of *Atriplex* and *Chenopodium* as well as *C. vulvaria* (Mandák et al., 2018), and the allotetraploidisation event leading to quinoa (Jarvis et al., 2017). We detected ChenSat-1a in all *Chenopodium* species tested, as well as in *Atriplex hortensis*, indicating presence of ChenSat-1a in the common ancestor of *Chenopodium* and *Atriplex*. Similarly, as members of the ChenSat-2 superfamily have been detected in all *Chenopodium* and *Atriplex* species tested, we suggest presence of a common ChenSat-2 progenitor family, most likely similar to ChenSat-2b or at least closely related. The ChenSat-2b repeat persisted in A genomes, but has been reduced in B genomes. In addition, ChenSat-2e emerged in A genome diploids, whereas ChenSat-2c and ChenSat-2d evolved in B genomes. ChenSat-2a is most likely the youngest family, as it was absent in most A- and B-containing species, except *C. pallidicaule*. We can therefore exclude *C. pallidicaule* as parental species for *C. quinoa*. Moreover, ChenSat-1b evolved in B genomes, presumably from divergence of ChenSat-1a monomers. Dating back at least 3 million years, allotetraploidisation of maternal A and paternal B genomes has led to *C. quinoa* (Kolano et al., 2016), containing ChenSat-1a, ChenSat-1b, and ChenSat-2b to ChenSat-2e tandem repeats. For the rDNAs, after *Chenopodium* speciation, the 5S and 18S-5.8S-25S rDNA spacers began to diverge and form A- and B-specific *Chenopodium* variants. The two A- and two B-derived 5S rDNA major sites are added to generate four 5S rDNA major sites in *C. quinoa*. In contrast, only B-derived 18S-5.8-25S rDNA is detectable in *C. quinoa* (Maughan et al., 2006, Kolano et al., 2016), whereas the A variant has been lost.

For the ChenSat-1 family, we present evidence that the subfamilies ChenSat-1a and ChenSat-1b are related. First, both satDNAs have short monomer sizes below 50 bp and about 60 % sequence similarity. Second, they occur close to each other, as detected on long reads and in the reference genome assembly, and also confirmed by multi-colour FISH with ChenSat-1a and ChenSat-1b probes. As ChenSat-1a occurs ubiquitously in *Chenopodium* species, whereas ChenSat-1b is restricted to B genomes, we assume that the B-specific ChenSat-1b emerged from ChenSat-1a by accumulation of mutations in a B genome precursor. The increase in monomer size may have occurred by replication slippage, as has been suggested for microsatellites (Viguera et al., 2001), by unequal crossing over, or by repair of double strand breaks. ChenSat-1b emerged without replacement of ChenSat-1a, indicating incomplete homogenisation. This is striking, as newly emerging families often replace the progenitor (Dover, 1982, Plohl et al., 2012). This may point to a structurally important role of ChenSat-1a, potentially forming the active centromere.

For the ChenSat-2 family, we observed diversification, with at least five ChenSat-2-derived subfamilies in different A and B genomes of *Chenopodium*. All ChenSat-2 subfamilies are most likely derived from a presumed common progenitor. Among other hallmarks, we observed a highly conserved monomer size (170-171 bp) for all ChenSat-2 subfamilies, likely important for DNA phasing (Melters et al., 2013), and an indication of selectional constraints. We used several approaches to corroborate the different abundance patterns and genome-specificities, such as read mapping, quantification in the assembled *C. quinoa* subgenomes, Southern hybridisation, and FISH to allotetraploid *C. quinoa*. The restriction to individual (sub)genomes indicates a rapid and species-specific ChenSat-2 evolution leading to a variety of different subfamilies. Similar observations have been reported for various plants and animals (Kopecna *et al*., 2012, Cai *et al*., 2014, Liu *et al*., 2019):

As ChenSat-2b is most widespread with presence in distantly related *C. vulvaria*, *A. hortensis*, and even *L. polyspermum*, we suggest ChenSat-2b or a precursor ChenSat-2b variant as the progenitor sequence. However, reduced ChenSat-2b signals in B genome diploids indicates an incomplete elimination from these species, explainable for example by molecular drive (Dover, 1982, Dover, 2002).

ChenSat-2a is exclusively present in *C. pallidicaule*, but absent in other A-containing genomes such as *C. quinoa*, explainable by two scenarios: (1) ChenSat-2a may have emerged after *C. pallidicaule* speciation, thus effectively excluding *C. pallidicaule* as potential parent of *C. quinoa*. This is consistent with data from genome sequencing considering *C. pallidicaule* as an unlikely progenitor of *C. quinoa* (Jarvis et al., 2017). (2) Alternatively, ChenSat-2a may have been eliminated from other A-containing *Chenopodium* genomes analysed here. This has been observed for example in natural and synthetic *Nicotiana tabacum* allopolyploids, in which continuous *Nic*CL3 satDNA arrays specific for the diploid progenitor *N. tomentosiformis* have been lost (Renny-Byfield et al., 2012). Nevertheless, multiple losses are necessary to explain the observed ChenSat-2a distribution. Therefore, we consider the emergence of ChenSat-2a in *C. pallidicaule* as more likely scenario.

We did not observe the rise of new satDNA families in allopolyploid quinoa, as documented in other polyploids. In *Nicotiana* allopolyploids which are older than five million years (*N. nesophila*, *N. stocktonii*, and *N. repanda*), new satDNAs have evolved, and sometimes have amplified to replace the parental satDNAs (Koukalova et al., 2010).

### Tandem repeats may provide targets for recombination between homoeologous chromosomes in allopolyploid quinoa

With two distinct repeat landscapes of the A- and B-derived subgenomes, quinoa is well-suited to investigate the invasion of satDNA into the respective other subgenome. In plants with a similar genome composition such as *Nicotiana* allopolyploids older than one million years (*N. quadrivalvis* and *N. clevelandii*), an exchange of satellite sequences between homoeologous chromosomes was already suspected (Koukalova et al., 2010). In the allopolyploid *C. quinoa* genome, recombination between homoeologous chromosomes are assumed to be rare, however some incidents were detected already two decades ago (Ward, 2000). Using the most current reference genome assembly, only a small number of homoeologous gene pairs (3.1%) has been mapped within the same subgenome, suggesting that recombination and chromosomal rearrangements have occurred between the A and B subgenomes to a small extent (Jarvis et al., 2017). Accordingly, we mapped A- and B-derived satDNAs to the same scaffold in only six cases (Table S3), possibly indicating interlocus recombination. However, using single molecule long reads originating from a single genomic region, we did not identify co-localisation of A-specific and B-specific *Chenopodium* rDNA variants. Nevertheless, we provide evidence that short ChenSat-2e arrays, enriched in the A genome, co-occur with the B-derived ChenSat-2c satDNA family, and confirm co-localisation on the same chromosome by multi-colour FISH and long read data.

Taken together, using short and long read bioinformatics as well as Southern and fluorescent *in situ* hybridisation, we traced seven satDNAs through *Chenopodium* speciation and allopolyploidisation. We observed satDNA diversification, replacement, reduction, and identified repeat families highly amplified in either the A or B genome diploids. After re-unification of both genomes in the allopolyploid quinoa, four of the seven satDNAs were subgenome-specific. We observed intermingling of satDNA families, which may point to homoeologous exchange of the ChenSat sequences. However, for the 5S rRNA genes, we can confidently suggest a strict separation of sequences on the A and B subgenomes.

## EXPERIMENTAL PROCEDURES

### Read clustering, tandem repeat identification and generation of representative monomeric consensus sequences

To identify satDNA from tetraploid *C. quinoa* and its diploid relatives, we used *RepeatExplorer* in comparative mode (Novák et al., 2010, Novák et al., 2013). We analysed the reads from the quinoa genome projects (Yasui et al., 2016, Jarvis et al., 2017) deposited at the NCBI sequence read archive: *Chenopodium pallidicaule* (SRR4425239), *Chenopodium suecicum* (SRR4425238), and *Chenopodium quinoa* (DRR057249). Read pre-treatment and interlacing was performed with custom scripts accompanying the local *RepeatExplorer* installation (paired_fastq_filtering.R and fasta_interlacer followed by seqclust). The reads were quality-trimmed to include only sequences with a Phred score ≥ 10 over 95 % of the read length. Overlapping read pairs have been excluded. Before comparative clustering, we randomly sampled 1.7 million paired shotgun pre-treated reads for *C. pallidicaule*, *C. suecicum*, and *C. quinoa* each, from which *RepeatExplorer* automatically chose 1,265,058, 1,263,518, and 1,265,808 reads, respectively. The resulting clusters have been classified by similarity searches against the Conserved Domain Database for the functional annotation of proteins (Marchler-Bauer et al., 2011), RepBase Update (Jurka et al., 2005), the REXdb database (Neumann et al., 2019), and a custom library containing common plant sequences (e.g. ribosomal, telomeric and plastid sequences). Clusters connected by mates and with matching annotations have been combined manually to superclusters.

Clusters with satellite-typical star-like and circular graph representations (Novák et al., 2010) were selected for further analysis. The *RepeatExplorer*-derived contigs were assembled and putative monomers were detected using *Tandem Repeats Finder* (Benson, 1999). In order to derive species-specific consensus sequences, iterative mapping of 2×1.7 million paired random short reads has been conducted for each genome until the consensus remained stable. Comparative quantification (Figure 3) relied on this read mapping dataset. The monomer consensus sequences are available in Data S1.

### Sequence comparison

Multiple sequence alignments were generated with the *MAFFT* (Katoh and Standley, 2013) and *MUSCLE* (Edgar, 2004) local alignment tools. They have been manually refined and used for the calculation of pairwise sequence identities with *MEGA X* (Kumar et al., 2018). We explored and visualised sequences with the multi-purpose software *Geneious 6.1.8* (Kearse et al., 2012). Dotplots have been generated with *FlexiDot* (Seibt et al., 2018) with wordsizes as indicated in the respective figure legends.

### Computational localisation of tandem repeats along the *C. quinoa* scaffolds

Tandem repeat positions on the *C. quinoa* pseudochromosomes and scaffolds (Jarvis et al., 2017) have been deduced by local *BLASTn* of a tandem repeat dimer. We retained only hits with an e-value ≤ 10^−10^, and transferred the hits into gff3 format. For scaffolds, we counted the number of hits for the specific satDNA (sub)family and deduced the scaffold’s origin from the A or B subgenome.

### Detection of higher order arrangements and interlocus homogenisation

We analysed higher order arrangements of tandem repeats on available *C. quinoa* single molecule real-time (SMRT) reads from accession number DRR057268. To account for the sequence error of the long reads, we used a nucleotide Hidden Markov Model (nHMM)-based approach. For each tandem repeat, we generated an nHMM from Illumina reads mapped to the respective consensus and used *nhmmer* (Wheeler and Eddy, 2013) to infer monomers along the SMRT reads. Hits were filtered individually for each satDNA family with parameters indicated in Table S2. Local monomer organisation was analysed visually using *FlexiDot* self dotplots (Seibt et al., 2018).

For detection of 5S rDNA interlocus homogenisation, individual monomers have been retrieved from the respective SMRT reads with the highest monomer count. The monomers have been aligned by *MAFFT* (Katoh and Standley, 2013), grouped by Neighbor Joining analysis and visualised with the *ETE toolkit* library (Huerta-Cepas et al., 2010).

### Plant material, DNA isolation, polymerase chain reaction (PCR) and cloning

Plant material has been obtained from sources indicated in Table 2. The plants were grown in the greenhouse under long day conditions. We isolated DNA as described (Arumuganathan and Earle, 1991) using 2× CTAB buffer and the additive polyvinylpyrrolidone (PVP). Especially for *C. quinoa* DNA, it was essential to retrieve the DNA immediately after isopropanol precipitation without centrifugation to avoid contamination with metabolites.

From the *C. quinoa* reference monomers, outward facing primers have been designed (Table S4). For the amplification of satellite DNA probes for Southern hybridisation and FISH, PCR was carried out with the specific primer pairs. PCR reactions with 50 ng plasmid template were performed in 50 µl volume containing 10x DreamTaq buffer and 2.5 units of DreamTaq polymerase (Promega). Standard PCR conditions were 94 °C for 5 min, followed by 35 cycles of 94 °C for 1 min, primer-specific annealing temperature for 30 sec, 72 °C for 1 min and a final incubation time at 72 °C for 5 min. We cloned and sequenced the PCR products for each satDNA family, and selected the one with the highest identity to the reference monomer as probe for subsequent hybridisations. We deposited sequences of satellite hybridisation probes online at the European Nucleotide Archive under the accessions LR215734 to LR215739 (study: http://www.ebi.ac.uk/ena/data/view/PRJEB31131).

### Southern hybridisation

For comparative Southern blots, restriction enzymes specific for each tandem repeat have been selected based on bioinformatics and practical tests of different enzymes. Genomic DNA was restricted, separated on 2 % agarose gels and transferred onto Hybond-N+ nylon membranes (GE Healthcare) by alkaline transfer. Hybridisations were performed according to standard protocols using probes labelled with ^32^P by random priming (Sambrook et al., 1989). Filters were hybridised at 60 °C and washed at 60 °C for 10 min in 2× SSC/ 0.1 % SDS. Signals were detected by autoradiography.

### Probe labelling, metaphase preparation, genomic and fluorescent *in situ* hybridisation

The satellite-specific probes were labelled by PCR in the presence of biotin-16-dUTP (Roche). We used nick translation to mark the probes for the ribosomal genes. The probe pZR18S containing a 5066 bp fragment of the sugar beet 18S-5.8S-25S rRNA gene (HE578879, Paesold et al., 2012) was labelled with DY-415 or DY-647-dUTP (Dyomics), whereas the probe pXV1 (Schmidt et al., 1994) for the 5S rRNA gene was labelled with digoxygenin-11-dUTP. For GISH with B-subgenome-specific probes, genomic DNA of *C. suecicum* has been heated to 99 °C for 10 min prior to labelling with digoxygenin-11-dUTP, also by nick translation.

We prepared mitotic chromosomes from the meristem of young leaves. Prior to fixation in methanol:glacial acetic acid (3:1), leaves were incubated for 2.5 to 3 h in 2 mM 8-hydroxyquinoline. Fixed plant material was digested for 4.5 hours at 37° C in an enzyme mixture consisting of 4 % (w/v) cellulase Onozuka R10 (Sigma 16419) and 20 % (v/v) pectinase from *Aspergillus niger* (Sigma P4716) in citrate buffer (4 mM citric acid and 6 mM sodium citrate) according to Kolano et al. (2011). After maceration, the mix was incubated for additional 25 min, before chromosome spreading by dropping according to Schwarzacher and Heslop-Harrison (2000) with modifications for beet (Schmidt et al., 1994).

We used FISH and GISH procedures described previously (Heslop-Harrison et al., 1991) with modifications for Amaranthaceae plants (Schmidt et al., 1994). Chromosome preparations were counterstained with DAPI (4’, 6’-diamidino-2-phenylindole) and mounted in antifade solution (CitiFluor). We examined the slides with a Zeiss Axioimager M1 UV epifluorescence microscope with appropriate filters, and equipped with an ASI BV300-20A camera coupled with the Applied Spectral Imaging software. The images were processed with Adobe Photoshop C5 software (Adobe Systems, San Jose, CA, USA) using only contrast optimisation, Gaussian and channel overlay functions affecting the whole image equally.

## ACKNOWLEDGEMENTS

We dedicate this paper to Ines Walter and Prof. Dr. Thomas Schmidt, who tragically and unforeseeably passed away during the course of this work. We thank Susan Liedtke’s excellent technical assistance with quinoa *in situ* hybridisations, Janine Hoffmann for assistance with cloning, and Falk Zakrzewski for an initial *RepeatExplorer* run of *C. quinoa* reads. We thank Kathrin M. Seibt for critical reading of the manuscript. The TU Dresden Center for Information Services and High Performance Computing (ZIH) is gratefully acknowledged for computer time allocations. Furthermore, we acknowledge Prof. Luz Gomez-Pando of the Universidad Nacional Agraria La Molina, the genebank at the IPK Gatersleben, and the U.S. National Plant Germplasm System for providing plant seeds.

## AUTHOR CONTRIBUTIONS

TH wrote the paper, made the figures, performed the sequence analyses and contributed to study design. BW analysed the chromovirus and assigned it to the CRM clade. IW performed the Southern and *in situ* hybridisations, and encouraged a stronger focus on the polyploid genomes. CO initially identified and cloned the tandem repeat sequences, as well as selected enzymes suitable for restriction. TS contributed to the research design and paper writing.

## DISCLOSURE DECLARATION

The authors declare no conflict of interests.

## SHORT LEGENDS FOR SUPPORTING INFORMATION

**Appendix S1:** Inferring of subgenome-specificity by analysis of the *RepeatExplorer* graph representations.

**Figure S1:** Genome contribution of *C. pallidicaule*, *C. suecicum* and *C. quinoa* to eight read clusters containing the satDNAs ChenSat-1a to ChenSat-1b, and ChenSat-2a to ChenSat-2e.

**Figure S2:** Sequence similarities among the *Chenopodium* satDNA families ChenSat-2a to ChenSat-2e.

**Figure S3:** Alignment of *Chenopodium* and other plant 5S rDNA sequences.

**Figure S4**: Arrangement of *C. quinoa* satDNA and 5S rDNA monomers in arrays, higher order, or inversions on SMRT reads.

**Figure S5:** ChenSat-1a arrays are often interrupted by LTR retrotransposons of the CRM chromovirus clade.

**Figure S6:** The LTR retrotransposon embedded in ChenSat-1a belongs to the CRM-type retrotransposons.

**Table S1:** Number of *C. quinoa* SMRT reads with each tandem repeat family.

**Table S2:** Co-occurrence of tandem repeats on *C. quinoa* SMRT reads.

**Table S3:** Unassigned *C. quinoa* scaffolds from the study of Jarvis et al. (2017), which we can newly assign to either the A or B subgenome.

**Table S4:** Primer for the amplification of *Chenopodium* tandem repeats.

**Data S1:** Consensus sequences from tandem repeat monomers in fasta format.

