## Supplementary Material for "Satellite DNA landscapes after allotetraploidisation of quinoa (*Chenopodium quinoa*) reveal unique A and B subgenomes"

Supplementary Data for

### **Tandem repeat evolution during *Chenopodium* speciation and formation of the allotetraploid crop quinoa (*Chenopodium quinoa*)**

Tony Heitkam, Beatrice Weber, Ines Walter, Charlotte Ost, and Thomas Schmidt

Corresponding author: Tony Heitkam

**This PDF file includes:**

Appendix S1

Figures S1 to S6

Table S1 to S4

Data S1

References in the supplement

#### Appendix S1

#### Analysis of the *RepeatExplorer* read clusters to deduce the tandem repeat amplification in the analysed *Chenopodium* genomes

As the read cluster abundance (visualised in Fig. 1) can misrepresent the real satDNA contribution (Novák et al., 2010, Ruiz-Ruano et al., 2016), we analysed the graph representations of all eight satDNA clusters (Fig. S1).

The two satDNA families ChenSat-1a and ChenSat-1b are represented by single repeat clusters, each (Fig. S1A-B). All reads from these clusters correspond exclusively to ChenSat-1a or ChenSat-1b, respectively. Therefore, graph interpretation is straightforward: As the ChenSat-1a cluster contains *C. pallidicaule, C. suecicum,* and *C. quinoa* reads, ChenSat-1a occurs in all tested genomes. Similarly, as the ChenSat-1b cluster harbours *C. suecicum* and *C. quinoa* reads, ChenSat-1b is presumably B-specific.

In contrast, the remaining satDNAs are part of larger repetitive clusters (ChenSat-2a, ChenSat-2b, ChenSat-2e), fall into multiple clusters (ChenSat-2c), or both (ChenSat-2d). ChenSat-2a and ChenSat-2b do not contain only the satDNA-typical circular graphs, but also additional repeats. To infer subgenome-specificity, only the circular representations are considered (Fig. S1C-D, arrows). The circular graphs enclose reads exclusively from A-containing genomes, indicating A-genome-specificity.

ChenSat-2c forms two clusters, both containing only reads from B-containing genomes, pointing to ChenSat-2c's B-genome-specificity (Fig. S1E-F).

Only the marked circular representations (Fig. S1G-H, arrows) are representative for ChenSat-2d. As they harbour only reads from *C. suecicum* and *C. quinoa*, we presume B-genome-specificity.

In addition to ChenSat-2d, CL59 contains an additional globular domain, corresponding to ChenSat-2e (Fig. S1H, marked by arrowheads). As it contains only reads from *C. pallidicaule* and *C. quinoa,* we suggest that ChenSat-2e is highly amplified in A-containing genomes.

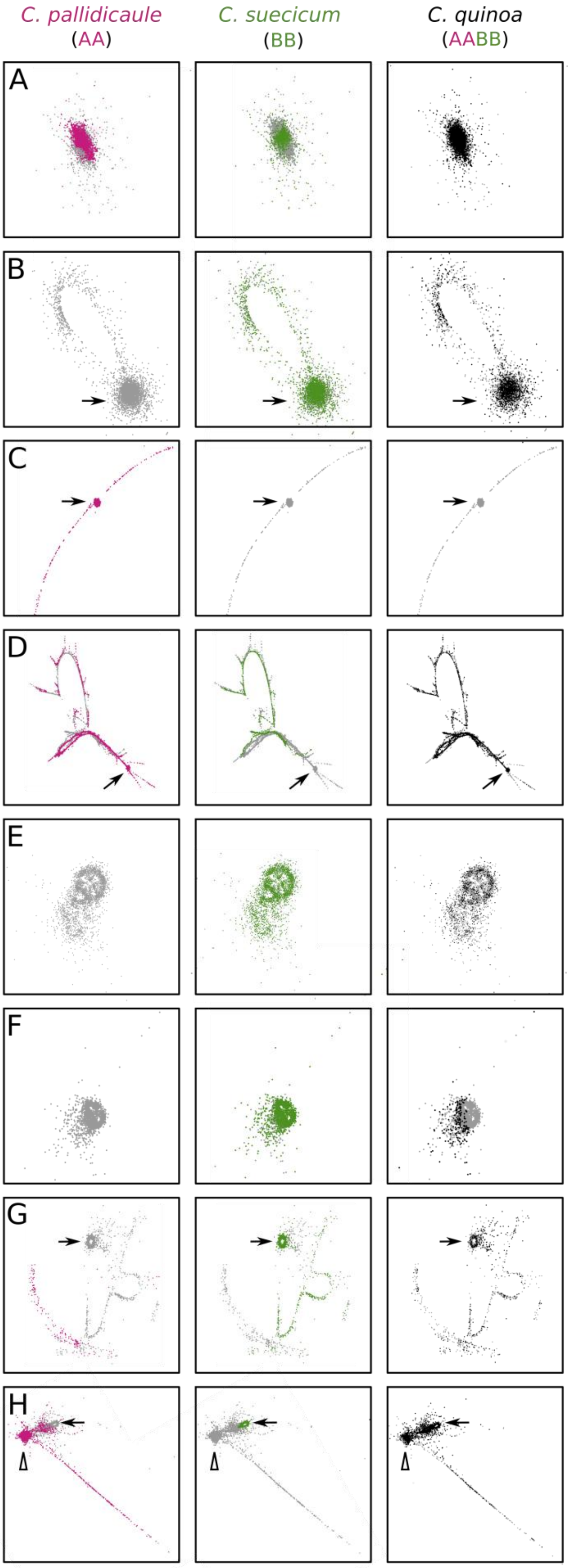


ChenSat-1a, CL6
all genomes contribute hits to globular structure

ChenSat-1b, CL102
only *C. pallidicaule* and *C. quinoa* genomes contribute hits to globular structure (arrows)

ChenSat-2a, CL123
only the *C. pallidicaule* genome contributes hits to globular structure (arrows)

ChenSat-2b, CL56
only *C. pallidicaule* and *C. quinoa* genomes contribute hits to globular structure (arrows)

ChenSat-2c, CL108
only *C. suecicum* and *C. quinoa* genomes contribute hits to globular structure

ChenSat-2c, CL141
only *C. suecicum* and *C. quinoa* genomes contribute hits to globular structure

ChenSat-2d, CL135
only *C. suecicum* and *C. quinoa* genomes contribute hits to globular structure (arrows)

ChenSat-2d, ChenSat-2e, CL59
only *C. suecicum* and *C. quinoa* genomes contribute hits to globular structure of ChenSat-2d (arrows)
only *C. pallidicaule* and *C. quinoa* genomes contribute hits to globular structure of ChenSat-2e (arrowheads)

**Fig. S1: Genome contribution of *C. pallidicaule*, *C. suecicum* and *C. quinoa* to eight read clusters containing the satDNAs ChenSat-1a to ChenSat-1b, and ChenSat-2a to ChenSat-2e.**

Each cluster is visualised three times with reads from each of the three analysed genomes marked in magenta (*C. pallidicaule*), green (*C. suecicum*), and black (*C. quinoa*). **(A)** ChenSat-1a: All three genomes contribute to the cluster, indicating ChenSat-1a occurrence in each genome. **(B)** ChenSat-1b: Only reads from *C. suecicum* and *C. quinoa* form the ChenSat-1b cluster, indicating B genome specificity of ChenSat-1b. **(C)** ChenSat-2a: The ChenSat-2a cluster is formed from *C. pallidicaule* reads only, indicating sole occurrence in the *C. pallidicaule* genome. **(D)** ChenSat-2b: This complex cluster harbours a tandem repeat part (arrow), corresponding to ChenSat-2b. It is only formed from *C. pallidicaule* and *C. quinoa* reads, indicating A-genome-specificity. **(E-F)** ChenSat-2c: This repeat is subdivided into two clusters (CL108 and CL141). Only reads from *C. suecicum* and *C. quinoa* form these clusters, indicating B-genome-specificity of ChenSat-2c. **(G)** ChenSat-2d: CL135 is a complex repeat cluster with the ChenSat-2d tandem repeat marked by arrows. Only *C. suecicum* and *C. quinoa* contribute to the ChenSat-2d tandem repeat, indicative of B-genome-specificity. **(H)** CL59 contains two genome-specific globular structures marked by arrows (ChenSat-2d) and arrowheads (ChenSat-2e).

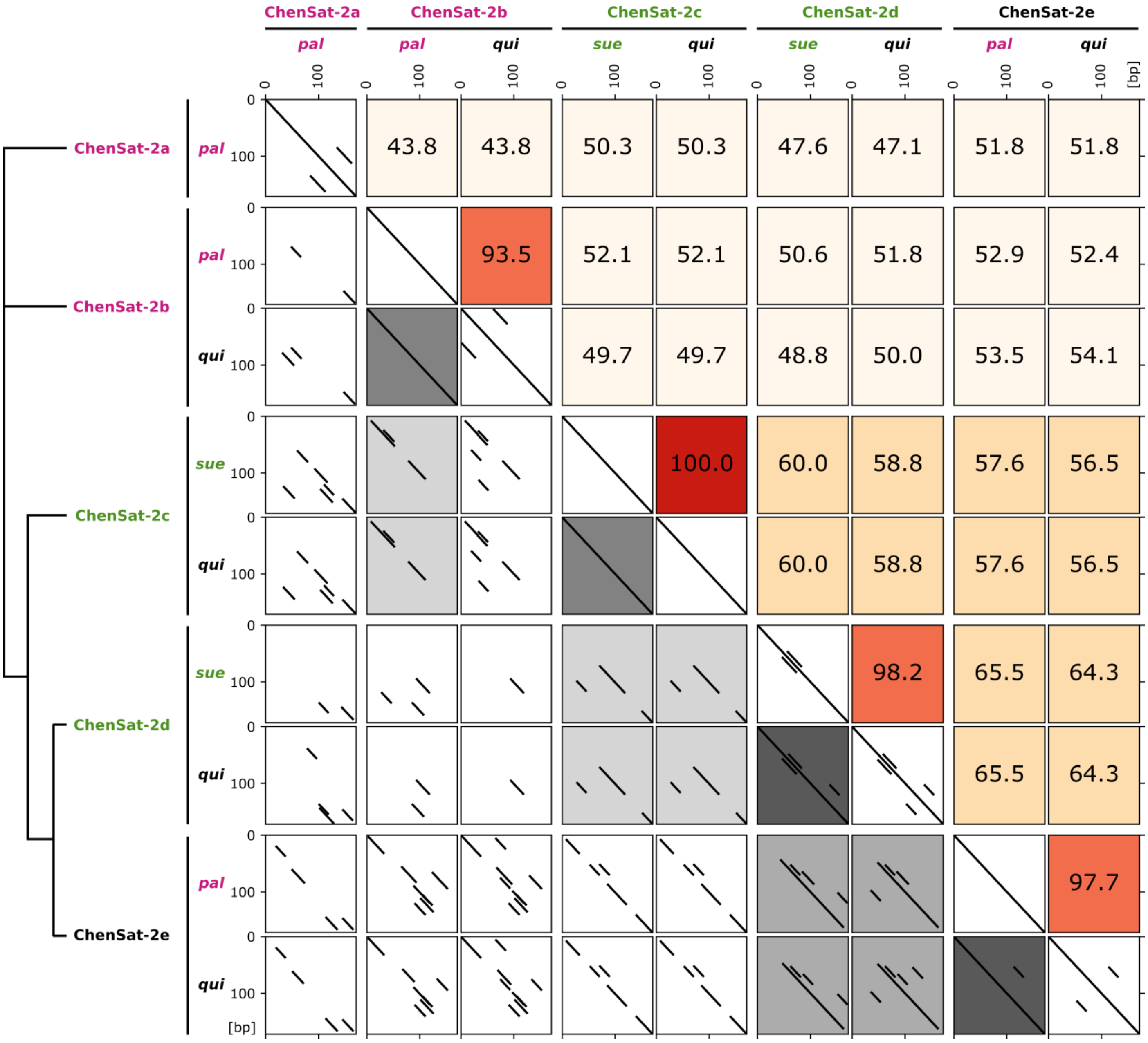


**Fig. S2: Sequence similarities among the *Chenopodium* satDNA families ChenSat-2a to ChenSat-2e.** The multi-dotplot of the nine ChenSat-2 consensus sequences (Fig. 2) shows pairwise sequence comparisons leading to 36 pairwise dotplots (lower left corner), 9 self dotplots (middle diagonal), and 36 identity values (upper right corner). The pairwise dotplots are shaded in grey according to the length of the longest shared diagonal (= longest common subsequence), whereas the pairwise identities are shaded orange and red according to their value. The Neighbor-Joining dendrogram (left) shows the relationship among the families. The branch lengths are not to scale.

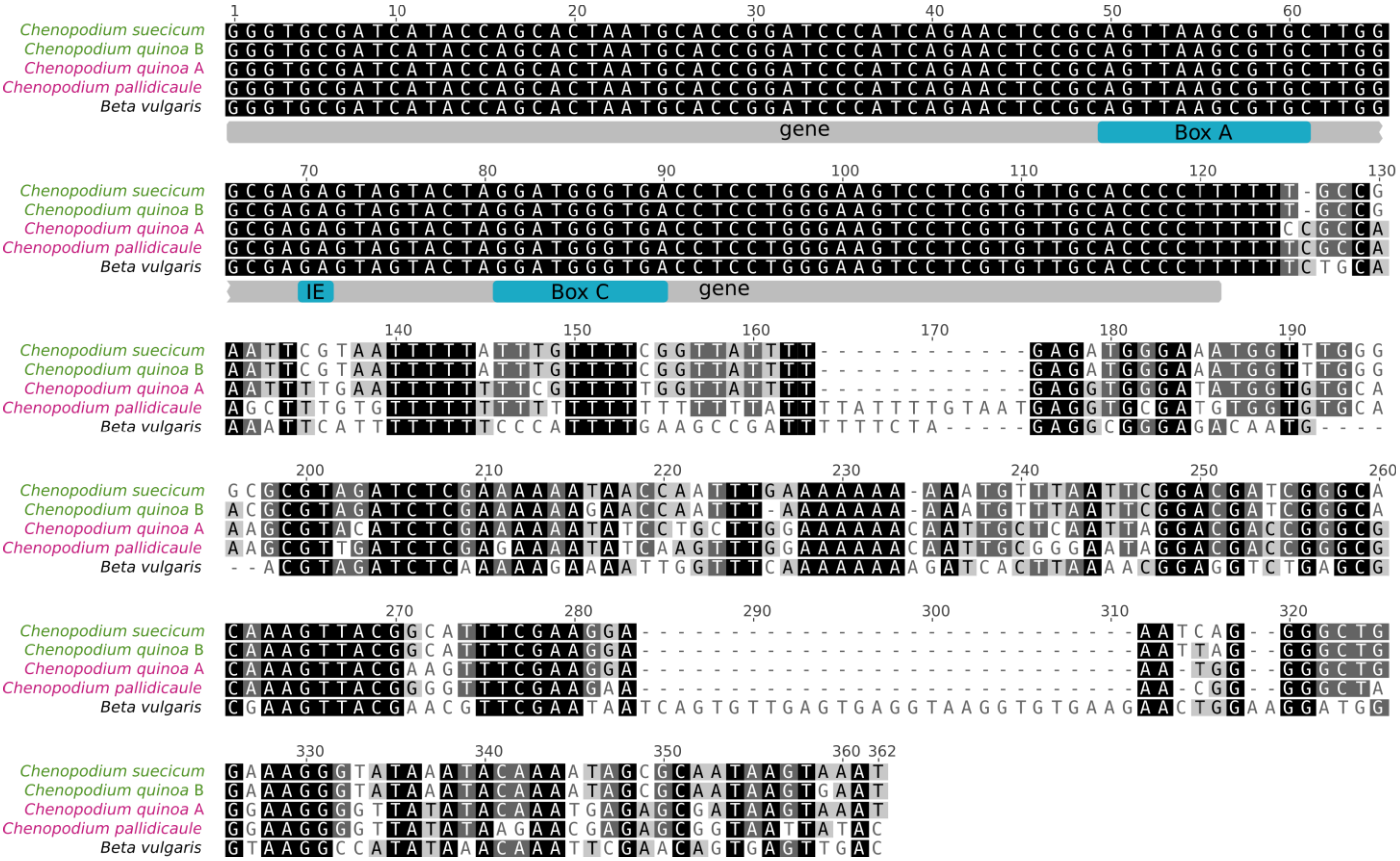


**Fig. S3: Alignment of *Chenopodium* and other plant 5S rDNA sequences.** Residues conserved in all compared sequences are shaded. The 5S rDNA monomeric consensus sequences of *C. pallidicaule*, *C. suecicum*, and both *C. quinoa* variants (derived from the A and B subgenome) have been aligned with a full monomer of *B. vulgaris* (Z25804). The coding region and the intergenic spacer were indicated. The region of the intragenic promoter is defined by three elements named Box A, Intermediate Element (IE), and Box C (Cloix et al., 2000), all marked in the alignment.

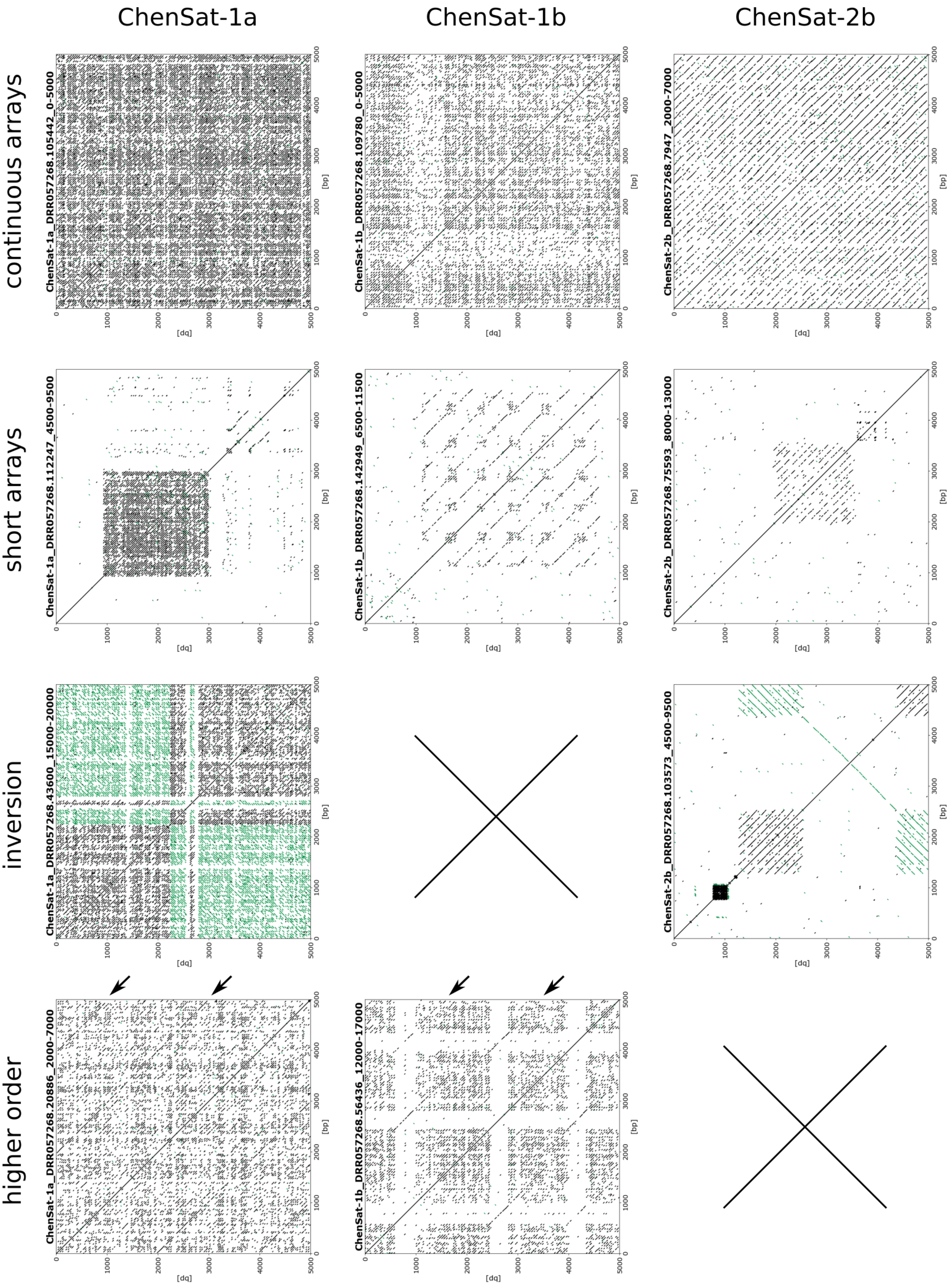

ChenSat-1a
ChenSat-1b
ChenSat-2b
continuous arrays
short arrays
inversion
higher order
ChenSat-1a_DRR057268.105442_0-5000
ChenSat-1b_DRR057268.109780_0-5000
ChenSat-2b_DRR057268.7947_2000-7000
ChenSat-1a_DRR057268.112247_4500-9500
ChenSat-1b_DRR057268.142949_6500-11500
ChenSat-2b_DRR057268.75593_8000-13000
ChenSat-1a_DRR057268.43600_15000-20000
ChenSat-2b_DRR057268.103573_4500-9500
ChenSat-1a_DRR057268.20886_2000-7000
ChenSat-1b_DRR057268.56436_12000-17000
[bp]
[dq]

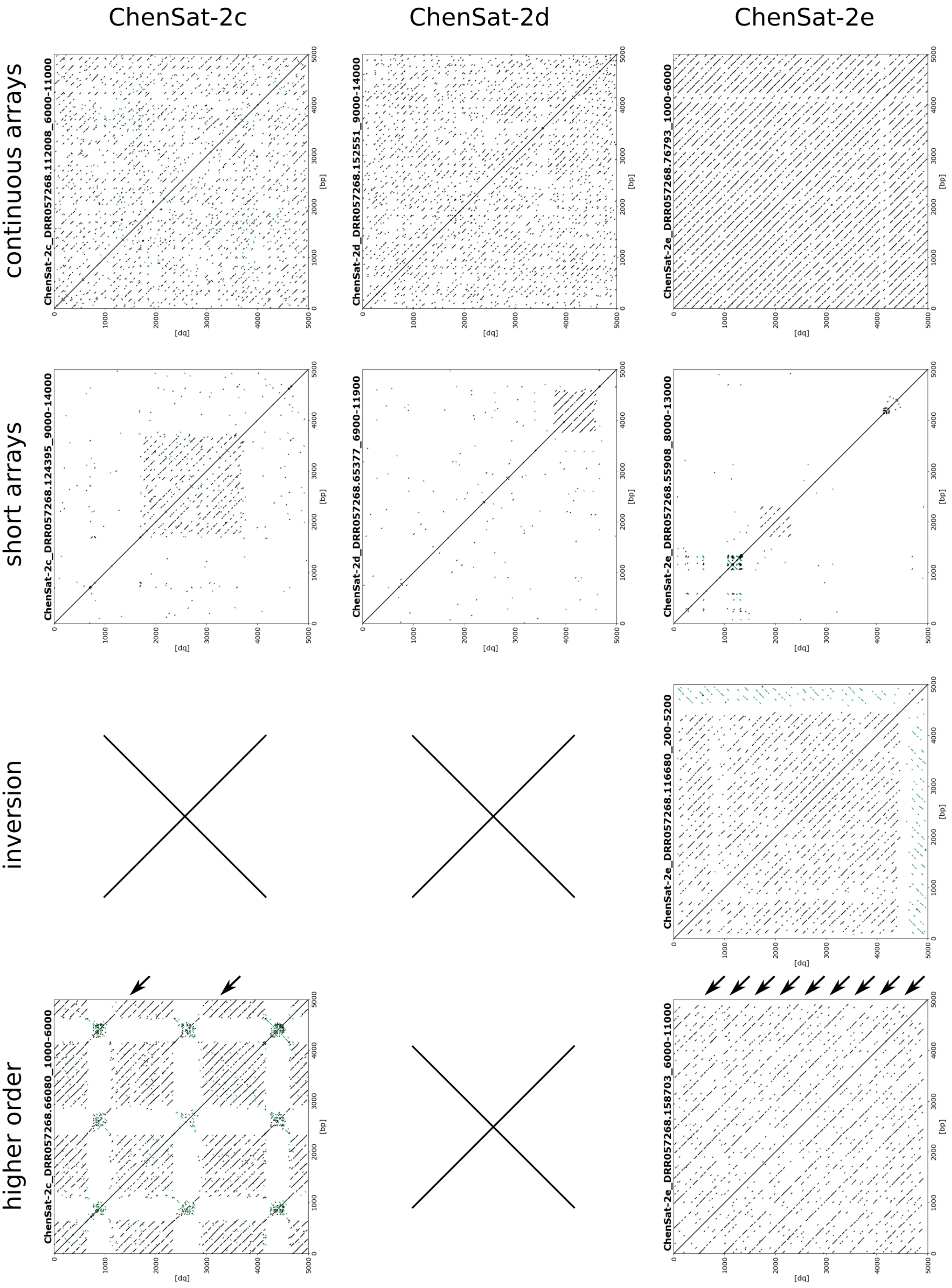
ChenSat-2c
ChenSat-2d
ChenSat-2e
continuous arrays
short arrays
inversion
higher order
ChenSat-2c_DRR057268.112008_6000-11000
ChenSat-2d_DRR057268.152551_9000-14000
ChenSat-2e_DRR057268.76793_1000-6000
ChenSat-2c_DRR057268.124395_9000-14000
ChenSat-2d_DRR057268.65377_6900-11900
ChenSat-2e_DRR057268.55908_8000-13000
ChenSat-2e_DRR057268.116680_200-5200
ChenSat-2c_DRR057268.66080_1000-6000
ChenSat-2e_DRR057268.158703_6000-11000
[dq]
[bp]

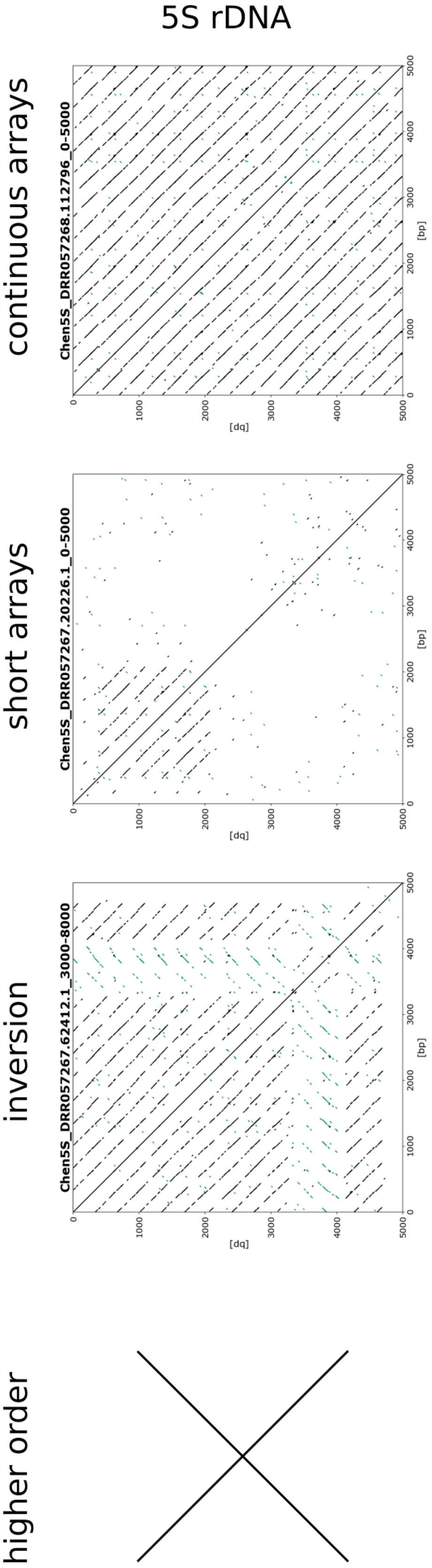


**Fig. S4**: **Arrangement of *C. quinoa* satDNA and 5S rDNA monomers in arrays, higher order, or inversions on SMRT reads.** We differentiate four arrangement patterns, i.e. continuous arrays, short arrays, higher order arrangements (arrows), and inversions. If detected, we show a self dotplot with a representative arrangement over a 5000 bp window. Forward matches are marked by black lines, whereas reverse matches are indicated in green. To account for the SMRT read error rate of 10-15 % (Rhoads and Au, 2015), we chose a wordsize of 20 bp and allowed five mismatches.

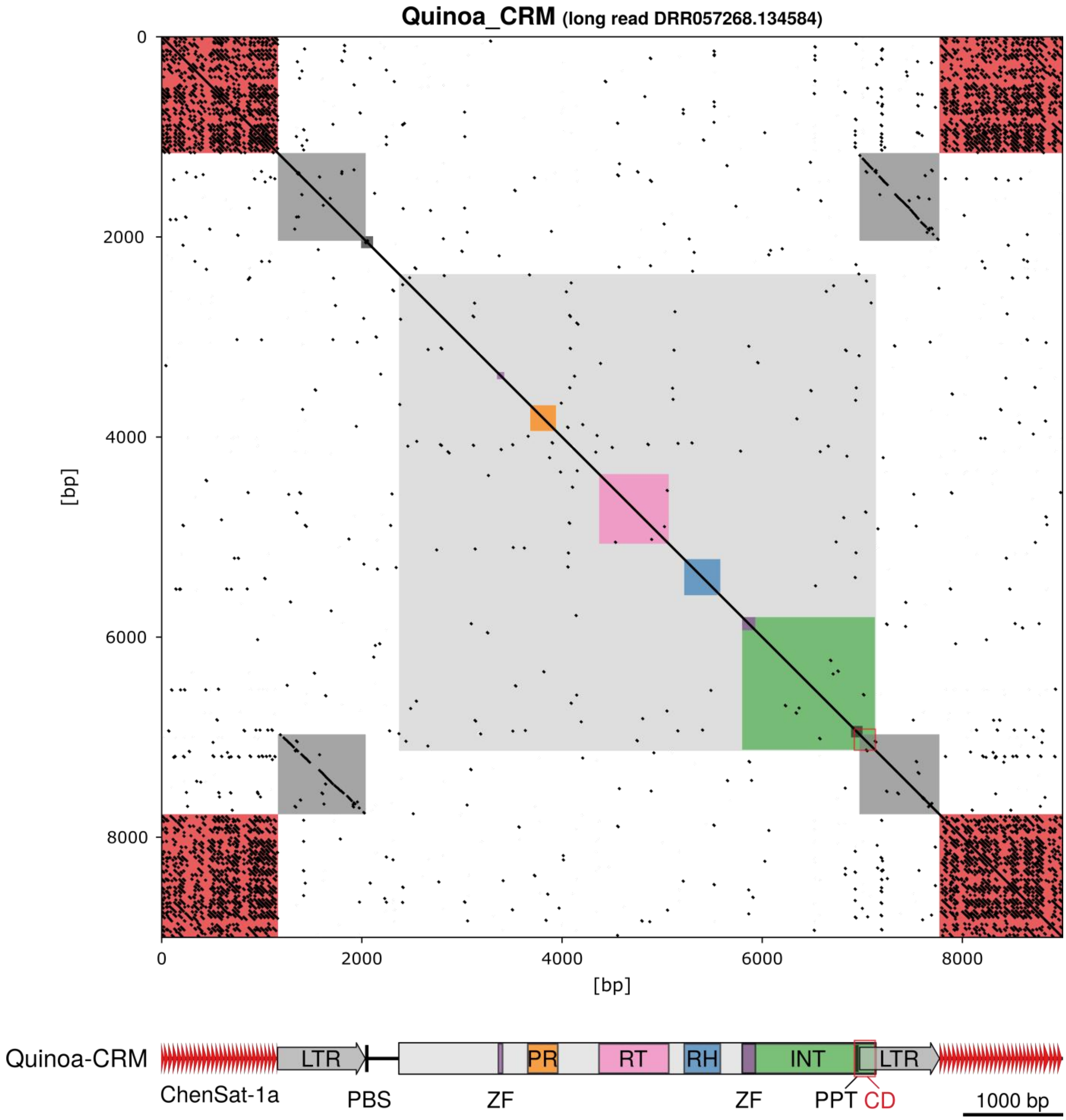


**Fig. S5: ChenSat-1a arrays are often interrupted by LTR retrotransposons of the CRM chromovirus clade. (top)** The dotplot of the *C. quinoa* SMRT read shows a typical ChenSat-1a satDNA array (red) with an LTR retrotransposon. The LTR is marked in dark grey. Structural features are colour-coded as indicated below. The dotplot was generated using a wordsize of 20 and tolerating five mismatches. **(bottom)** Structure of a representative *C. quinoa* CRM chromovirus, Quinoa-CRM, as detected interspersed in Chensat-1a. Open arrows represent the long terminal repeats (LTRs). Conserved domains are shown: The continuous ORF with the catalytic domains of protease (PR), reverse transcriptase (RT), RNaseH (RH), and integrase (INT), containing the chromodomain (CD) at the C-terminus. The ORF of the *gag-pol* polyprotein extends into the 3' LTR. Primer binding site (PBS), polypurine tract (PPT) and zinc finger (ZF) motifs are indicated.

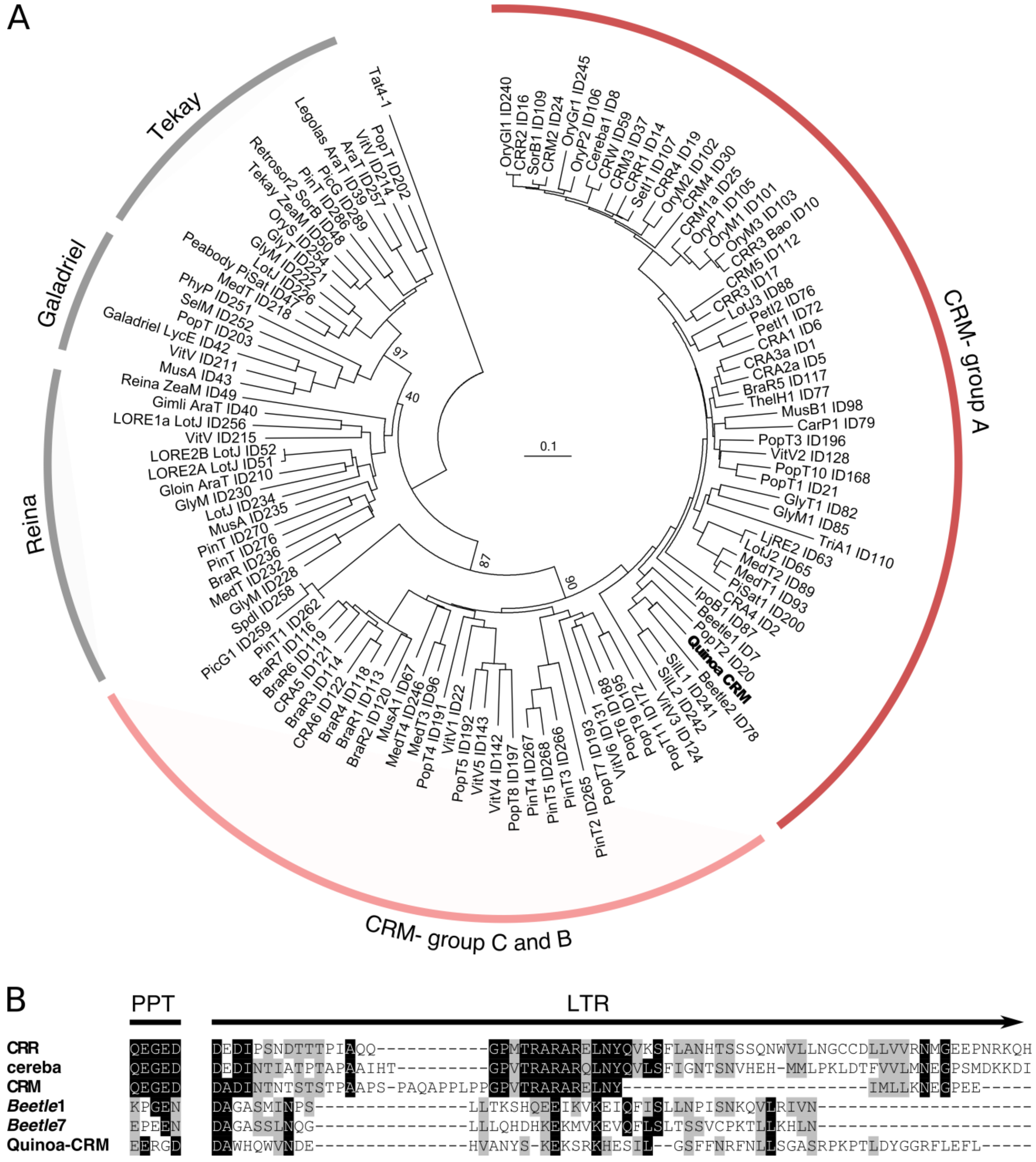


**Fig. S6: The LTR retrotransposon embedded in ChenSat-1a belongs to the CRM-type retrotransposons as indicated by its reverse transcriptase (RT) and chromodomain sequences.** For the Quinoa-CRM classification, we extracted a representative genomic copy from the high-quality quinoa genome assembly (Jarvis et al., 2017, *Cq_PI614886_v1*, Chr16: 32,720,303...32,726,561 bp). It is characterised by an intact, continuous open reading frame (ORF), a 5 bp target site duplication, and 100 % similar LTRs, indicative of a valid and recent transposition event. **(A)** A dendrogram of additional 116 chromovirus RTs (Neumann et al., 2011) shows the similarity to other centromeric chromoviruses such as *Beetle*1 from the related beet genome, clearly supporting the assignment to the CRM clade, group A. Tat4-1, a Ty3-*gypsy* retrotransposon of the sister lineage Ogre/Tat was chosen for rooting the phylogeny**.** **(B)** An alignment of the C-terminal chromodomain, embedded in the retrotransposon *pol* ORF, reveals the presence of conserved amino acids as described (Weber et al., 2013), indicative of a functional CRM-type chromodomain. As the *pol* ORF extends into the retrotransposon LTR, structural features, such as the position of the polypurine tract (PPT) and long terminal repeat (LTR) are marked. Columns with at least 50 % amino acid sequence identity/similarity are shaded in black/grey.

**Table S1:** Number of *C. quinoa* SMRT reads with each tandem repeat family and the nhmmer parameters to reproduce this result (Wheeler and Eddy, 2013).

| | **Minimum bitscore** | **Minimum nHMM coverage [bp]** | **SMRT read number** | **Total read count** |
|---|---|---|---|---|
| 5SA | 40 | 60 | 28 | **74** |
| 5SB | 40 | 60 | 46 | |
| ChenSat-1a | 15 | 30 | 5426 | **5653** |
| ChenSat-1b | 15 | 30 | 227 | |
| ChenSat-2a | 30 | 80 | 1 | **1592** |
| ChenSat-2b | 30 | 80 | 148 | |
| ChenSat-2c | 30 | 80 | 303 | |
| ChenSat-2d | 30 | 80 | 85 | |
| ChenSat-2e | 30 | 80 | 1055 | |
| **Total read count** | | | **7319** | |

**Table S2:** Co-occurrence of tandem repeats on *C. quinoa* SMRT reads. The number of reads with co-occurences is indicated. Co-occurences of A- and B-specific tandem repeats on a SMRT read are highlighted.

| Tandem repeats and genome specificity | | 5SA | 5SB | ChenSat-1a | ChenSat-1b | ChenSat-2a | ChenSat-2b | ChenSat-2c | ChenSat-2d | ChenSat-2e |
|---|---|---|---|---|---|---|---|---|---|---|
| | | A | B | --- | B | A | A | B | B | --- |
| 5SA | A | --- | --- | --- | --- | --- | --- | --- | --- | --- |
| 5SB | B | 0 | --- | --- | --- | --- | --- | --- | --- | --- |
| ChenSat-1a | --- | 0 | 2 | --- | --- | --- | --- | --- | --- | --- |
| ChenSat-1b | B | 0 | 0 | 6 | --- | --- | --- | --- | --- | --- |
| ChenSat-2a | A | 0 | 0 | 0 | 0 | --- | --- | --- | --- | --- |
| ChenSat-2b | A | 0 | 0 | 4 | 0 | 0 | --- | --- | --- | --- |
| ChenSat-2c | B | 0 | 0 | 3 | 1 | 0 | 0 | --- | --- | --- |
| ChenSat-2d | B | 0 | 0 | 2 | 1 | 0 | 0 | 0 | --- | --- |
| ChenSat-2e | --- | 1 | 0 | 18 | 1 | 0 | 65 | 91 | 6 | --- |

**Table S3:** Unassigned *C. quinoa* scaffolds from the study of Jarvis et al. (2017), which we can newly assign to either the A or B subgenome. For five scaffolds, we detected similarity to both A- and B-derived repeats, indicative of recombination (here labelled "conflict").

| Scaffolds Name | Size [bp] | ChenSat-1a | Highly abundant in A | A-specific repeats | | | B-specific repeats | | | | Number of newly assigned scaffolds |
|---|---|---|---|---|---|---|---|---|---|---|---|
| | | | ChenSat-2e | 5S rDNA spacer | ChenSat-2a | ChenSat-2b | 5S rDNA spacer | ChenSat-1b | ChenSat-2c | ChenSat-2d | |
| C_Quinoa_Scaffold_2387 | 1135038 | 0 | 16 | 398 | 0 | 6 | 0 | 0 | 0 | 0 | 24x: A |
| C_Quinoa_Scaffold_1747 | 7817241 | 0 | 17 | 106 | 0 | 8 | 0 | 0 | 0 | 0 | |
| C_Quinoa_Scaffold_1963 | 32410 | 0 | 0 | 102 | 0 | 0 | 0 | 0 | 0 | 0 | |
| C_Quinoa_Scaffold_1074 | 25841 | 0 | 0 | 82 | 0 | 0 | 0 | 0 | 0 | 0 | |
| C_Quinoa_Scaffold_1168 | 2263085 | 0 | 5 | 21 | 0 | 0 | 0 | 0 | 0 | 0 | |
| C_Quinoa_Scaffold_2127 | 11107391 | 8 | 41 | 1 | 0 | 26 | 0 | 0 | 0 | 0 | |
| C_Quinoa_Scaffold_4471 | 5387144 | 0 | 182 | 0 | 0 | 871 | 0 | 0 | 0 | 0 | |
| C_Quinoa_Scaffold_2356 | 51633 | 0 | 0 | 0 | 0 | 311 | 0 | 0 | 0 | 0 | |
| C_Quinoa_Scaffold_1628 | 49290 | 0 | 0 | 0 | 0 | 292 | 0 | 0 | 0 | 0 | |
| C_Quinoa_Scaffold_1083 | 3637715 | 16 | 2 | 0 | 0 | 250 | 0 | 0 | 0 | 0 | |
| C_Quinoa_Scaffold_2185 | 5863117 | 10 | 37 | 0 | 0 | 233 | 0 | 0 | 0 | 0 | |
| C_Quinoa_Scaffold_1695 | 3008951 | 8 | 1 | 0 | 0 | 186 | 0 | 0 | 0 | 0 | |
| C_Quinoa_Scaffold_1558 | 36924 | 0 | 0 | 0 | 0 | 176 | 0 | 0 | 0 | 0 | |
| C_Quinoa_Scaffold_3485 | 438987 | 562 | 18 | 0 | 0 | 123 | 0 | 0 | 0 | 0 | |
| C_Quinoa_Scaffold_2586 | 24844 | 0 | 0 | 0 | 0 | 89 | 0 | 0 | 0 | 0 | |
| C_Quinoa_Scaffold_3163 | 10083965 | 1483 | 51 | 0 | 0 | 35 | 0 | 0 | 0 | 0 | |
| C_Quinoa_Scaffold_4365 | 1955234 | 0 | 0 | 0 | 0 | 35 | 0 | 0 | 0 | 0 | |
| C_Quinoa_Scaffold_3500 | 7098671 | 0 | 101 | 0 | 0 | 20 | 0 | 0 | 0 | 0 | |
| C_Quinoa_Scaffold_2299 | 4034256 | 19 | 25 | 0 | 0 | 20 | 0 | 0 | 0 | 0 | |
| C_Quinoa_Scaffold_2588 | 8811694 | 357 | 62 | 0 | 0 | 14 | 0 | 0 | 0 | 0 | |
| C_Quinoa_Scaffold_2889 | 4265787 | 339 | 7 | 0 | 0 | 14 | 0 | 0 | 0 | 0 | |
| C_Quinoa_Scaffold_2674 | 1939591 | 0 | 16 | 0 | 0 | 13 | 0 | 0 | 0 | 0 | |
| C_Quinoa_Scaffold_4446 | 6153240 | 0 | 25 | 0 | 0 | 12 | 0 | 0 | 0 | 6 | |
| C_Quinoa_Scaffold_1559 | 7254884 | 3166 | 531 | 0 | 0 | 12 | 0 | 0 | 0 | 0 | |
| C_Quinoa_Scaffold_2033 | 94043 | 0 | 0 | 0 | 0 | 0 | 0 | 966 | 0 | 0 | 27x: B |
| C_Quinoa_Scaffold_2265 | 170979 | 0 | 0 | 0 | 0 | 0 | 0 | 872 | 0 | 0 | |
| C_Quinoa_Scaffold_1372 | 30275 | 0 | 0 | 0 | 0 | 0 | 0 | 527 | 0 | 0 | |
| C_Quinoa_Scaffold_2262 | 820779 | 292 | 0 | 0 | 0 | 0 | 0 | 500 | 123 | 0 | |
| C_Quinoa_Scaffold_2124 | 22138 | 0 | 0 | 0 | 0 | 0 | 0 | 305 | 0 | 0 | |
| C_Quinoa_Scaffold_1905 | 41012 | 0 | 0 | 0 | 0 | 0 | 0 | 249 | 0 | 0 | |
| C_Quinoa_Scaffold_3752 | 28736 | 0 | 0 | 0 | 0 | 0 | 0 | 245 | 0 | 0 | |
| C_Quinoa_Scaffold_1755 | 3648046 | 0 | 0 | 0 | 0 | 0 | 0 | 229 | 0 | 0 | |
| C_Quinoa_Scaffold_3146 | 79956 | 0 | 0 | 0 | 0 | 0 | 0 | 0 | 478 | 0 | |
| C_Quinoa_Scaffold_2539 | 585471 | 39 | 0 | 0 | 0 | 0 | 0 | 0 | 214 | 0 | |
| C_Quinoa_Scaffold_2865 | 2351721 | 0 | 0 | 0 | 0 | 0 | 0 | 0 | 181 | 0 | |
| C_Quinoa_Scaffold_1478 | 47161 | 0 | 0 | 0 | 0 | 0 | 0 | 0 | 172 | 0 | |
| C_Quinoa_Scaffold_2937 | 26498 | 0 | 0 | 0 | 0 | 0 | 0 | 0 | 126 | 0 | |
| C_Quinoa_Scaffold_1779 | 35590 | 0 | 0 | 0 | 0 | 0 | 0 | 0 | 51 | 0 | |
| C_Quinoa_Scaffold_3026 | 62613 | 0 | 0 | 0 | 0 | 0 | 0 | 0 | 50 | 0 | |
| C_Quinoa_Scaffold_1116 | 33452 | 0 | 0 | 0 | 0 | 0 | 0 | 0 | 48 | 0 | |
| C_Quinoa_Scaffold_3796 | 8186616 | 405 | 68 | 0 | 0 | 0 | 0 | 0 | 39 | 2 | |
| C_Quinoa_Scaffold_3387 | 843854 | 0 | 0 | 0 | 0 | 0 | 0 | 0 | 35 | 398 | |
| C_Quinoa_Scaffold_2756 | 28172 | 0 | 0 | 0 | 0 | 0 | 0 | 0 | 24 | 0 | |
| C_Quinoa_Scaffold_1088 | 2496132 | 2344 | 0 | 0 | 0 | 0 | 0 | 0 | 20 | 0 | |
| C_Quinoa_Scaffold_3850 | 3570316 | 1789 | 0 | 0 | 0 | 0 | 0 | 0 | 19 | 7 | |
| C_Quinoa_Scaffold_2368 | 1988798 | 0 | 0 | 0 | 0 | 0 | 0 | 0 | 19 | 0 | |
| C_Quinoa_Scaffold_3315 | 23411 | 0 | 0 | 0 | 0 | 0 | 0 | 0 | 17 | 0 | |
| C_Quinoa_Scaffold_1114 | 971808 | 3652 | 0 | 0 | 0 | 0 | 0 | 0 | 10 | 0 | |
| C_Quinoa_Scaffold_1334 | 1041137 | 0 | 0 | 0 | 0 | 0 | 0 | 0 | 10 | 0 | |
| C_Quinoa_Scaffold_3464 | 256749 | 0 | 0 | 0 | 0 | 0 | 0 | 0 | 0 | 372 | |
| C_Quinoa_Scaffold_2951 | 393229 | 0 | 0 | 0 | 0 | 0 | 0 | 0 | 0 | 10 | |
| C_Quinoa_Scaffold_3389 | 8125514 | 0 | 23 | 0 | 0 | 19 | 0 | 0 | 187 | 0 | 6x: Conflict |
| C_Quinoa_Scaffold_2048 | 11561360 | 9 | 71 | 0 | 0 | 14 | 0 | 2 | 0 | 13 | |
| C_Quinoa_Scaffold_1155 | 3984343 | 20 | 0 | 0 | 0 | 11 | 0 | 0 | 3 | 0 | |
| C_Quinoa_Scaffold_3035 | 12398513 | 588 | 67 | 0 | 0 | 8 | 0 | 0 | 8 | 18 | |
| C_Quinoa_Scaffold_2081 | 5763804 | 0 | 39 | 0 | 0 | 6 | 0 | 0 | 0 | 25 | |
| C_Quinoa_Scaffold_1034 | 3203711 | 571 | 1 | 0 | 0 | 5 | 0 | 0 | 0 | 16 | |

**Table S4:** Primer pairs for the amplification of *Chenopodium* tandem repeats.

| Probe | Primer | Sequence | Tm [° C] |
|---|---|---|---|
| ChenSat-1a | ChenSat1a-for | GTTTGACTTTCATTTGATTC | 49.1 |
| | ChenSat1a-rev | CGAAACAAACTTACACAA | 46.9 |
| ChenSat-1b | ChenSat1b-for | GGGCTCATTAGCCCTAAGGG | 61.4 |
| | ChenSat1b-rev | CCCCTTGGATGGGCGATG | 60.5 |
| ChenSat-2a | ChenSat2a-for | CCTTTCAAGTCATTTTTTGGGG | 56.5 |
| | ChenSat2a-rev | GGACTTTTTGAGCTAGAACTCG | 58.4 |
| ChenSat-2b | ChenSat2b-for | CAAGCATTGAGCTAAAATCATG | 54.7 |
| | ChenSat2b-rev | TGATTTAACTCCTATATTACACC | 53.5 |
| ChenSat-2c | ChenSat2c-for | AATGGCTACTCACTTGCC | 53.7 |
| | ChenSat2c-rev | CATTTCGGCTCAAAATGGC | 54.5 |
| ChenSat-2d | ChenSat2d-for | GCACATATGTCAATTTTCAACC | 54.7 |
| | ChenSat2d-rev | GCTTATTTCTCTTAATTTCTATCC | 54.2 |
| ChenSat-2e | ChenSat2e-for | GCCTATATCACTACTTGTATGCC | 58.9 |
| | ChenSat2e-rev | GCCATTTTGAATGAAATTGCGTCC | 59.3 |

**Data S1:** Fasta sequences of reference monomer sequences for the *Chenopodium* tandem repeats.

```
>ChenSat-1a_pal
AAAGCTAATTGAATCAAATGAAAGTCAAACACATTCAAAC
>ChenSat-1a_sue
AAAGCTTTTTGAATCAAATGAAAGTCAAACACATTCAAAC
>ChenSat-1a_qui
AAAGCTATTTGAATCAAATGAAAGTCAAACACATTCAAAC
```

```
>ChenSat-1b_sue
AAGGGGCTCATTAGCCCTAAGGGGCGTCAGACACATCATCGCCCATCC
>ChenSat-1b_qui
AAGGGGCTCATTAGCCCTAAGGGGCGTCAGACACATCATCGCCCATCC
```

```
>ChenSat-2a_pal
TGTTTGATGGTTTCATGGACAATAAAACACAATATAAACGAGTTCTAGCTCAAAAAGTCCAAAAATAGGCCTTTCAAGTCATTTTTTGGGGTTTA
CTTGCCTAATTCTTACATATCAATTCATTTCAACCTAGTTTTCACTTGTACATGCCTAAATCACATAGTTAATAG
```

```
>ChenSat-2b_pal
TATTTATGGCTTGAAACAATATAAAAAAAGCAAATTTACAAGTTTTAGGTTCATTTAGAGTATTTACGGTTCAAAATGGGCAAAATGACTTCTAA
CTTGCCCATTTTTCATGATTTTAGCTCAATGCTTGAATTAACTCCTATATTACACCTAAGATACTTAGTTATGAA
>ChenSat-2b_qui
TATTTATGGCTTGAAACAACCTAAAAAAAGCAATTTCACAAGTTTTAGGTTCATTTAGAGTGTTTACGGCTCGAAATGGCCTAAATGAGTTCTAA
CTTGCCCATTTTTCATGATTTTAGCTCAATGCTTGATTTAACTCCTATATTACACCTAAGATACTTAGTTATGAA
```

```
>ChenSat-2c_sue
TGTTTGTCATTTGAAATTACATAAAAATCGCCAAGTTCTAAGTTTTTGGGTCCAATTCATCAAAAAAAGCCATTTTGAGCCGAAATGGCTACTCA
CTTGCCTATTTTCTTCATTTCTCAACTTATAAACTCCTTGCATCTCAAAACAACTCATAATCACATATTTATAAG
>ChenSat-2c_qui
TGTTTGTCATTTGAAATTACATAAAAATCGCCAAGTTCTAAGTTTTTGGGTCCAATTCATCAAAAAAAGCCATTTTGAGCCGAAATGGCTACTCA
CTTGCCTATTTTCTTCATTTCTCAACTTATAAACTCCTTGCATCTCAAAACAACTCATAATCACATATTTATAAG
```

```
>ChenSat-2d_sue
TGCCTAAGGATAGAAATTAAGAGAAATAAGCACATATGTCAATTTTCGACCCTAACTCATCCAAAATCATCCAAATTTGGCCTATATGGCTAAAA
CCATGCCTATTTTCTATGTTTTCAAGCAAATGGACTCGTTCTTTCTTCTAACATTCCAATATCAAATACTTTTAAG
>ChenSat-2d_qui
TGCCTAAGGATAGAAATTAAGAGAAATAAGCACATATGTCAATTTTCGACCCTTACTCATCCAAAATCATCCAAATTTGGCCTATATGGCTAAAA
CCATGCCTATTTTCTATGTTTTCAATCAAATGGACTCGTTCTTTCTTCTAACATTCCAATATCAATTACTTTTAAG
```

```
>ChenSat-2e_pal
CAGCTAAGGCTTGAAATCACATTAAAAGAGCATTTGAATGGTTTTTGGACGCAATTTCATTCAAAATGGCCGAAATAAGGCCTATATCACTACTT
GTATGCCTATTTTCTATATTTTTAATCAAATTAGCTCATTCTTCCTCTTAACATGTCATTATTACCTAGTTAGAGG
>ChenSat-2e_qui
CAGCTAAGGCTTGAAATCACATTAAAAGAGCATTTGAAGGGTTTTAGGACGCAATTTCATTCAAAATGGCCCGAATAAGGCCTATATCACTACTT
GTATGCCTATTTTCTATATTTTTAATCAAATTAGCTCATTCTTCCTCTTAACATGTCATTATTACCTAGTTAGAGG
```

```
>5SrDNA_pal
GGGTGCGATCATACCAGCACTAATGCACCGGATCCCATCAGAACTCCGCAGTTAAGCGTGCTTGGGCGAGAGTAGTACTAGGATGGGTGACCTCC
TGGGAAGTCCTCGTGTTGCACCCCTTTTTTCGCCAAGCTTTGTGTTTTTTTTTTTTTTTTTTTTTATTTATTTTGTAATGAGGTGCGATGTGGT
GTGCAAAGCGTTGATCTCGAGAAAATATCAAGTTTGGAAAAAACAATTGCGGGAATAGGACGACCGGGCGCAAAGTTACGGGGTTTCGAAGAAAA
CGGGGGCTAGGAAGGGGTTATATAAGAACGAGAGCGGTAATTATAC
>5SrDNA_sue
GGGTGCGATCATACCAGCACTAATGCACCGGATCCCATCAGAACTCCGCAGTTAAGCGTGCTTGGGCGAGAGTAGTACTAGGATGGGTGACCTCC
TGGGAAGTCCTCGTGTTGCACCCCTTTTTTGCCGAATTCGTAATTTTTATTTGTTTTCGGTTATTTTGAGATGGGAAATGGTTTGGGGCGCGTAG
ATCTCGAAAAAATAACCAATTTGAAAAAAAAAATGTTTAATTCGGACGATCGGGCACAAAGTTACGGCATTTCGAAGGAAATCAGGGGCTGGAAA
GGGTATAAATACAAAATAGCGCAATAAGTAAAT
>5SrDNA_quiA
GGGTGCGATCATACCAGCACTAATGCACCGGATCCCATCAGAACTCCGCAGTTAAGCGTGCTTGGGCGAGAGTAGTACTAGGATGGGTGACCTCC
TGGGAAGTCCTCGTGTTGCACCCCTTTTTCCGCCAAATTTTGAATTTTTTTTCGTTTTTGGTTATTTTGAGGTGGGATATGGTGTGCAAAGCGTA
```

```
CATCTCGAAAAAATATCCTGCTTGGAAAAAACAATTGCTCAATTAGGACGACCGGGCGCAAAGTTACGAAGTTTCGAAGGAAATGGGGGCTGGGA
AGGGGTTATATACAAATGAGAGCGATAAGTAAAT
>5SrDNA_quiB
GGGTGCGATCATACCAGCACTAATGCACCGGATCCCATCAGAACTCCGCAGTTAAGCGTGCTTGGGCGAGAGTAGTACTAGGATGGGTGACCTCC
TGGGAAGTCCTCGTGTTGCACCCCTTTTTTGCCGAATTCGTAATTTTTATTTGTTTTCGGTTATTTTGAGATGGGAAATGGTTTGGGACGCGTAG
ATCTCGAAAAAAGAACCAATTTAAAAAAAAAAATGTTTAATTCGGACGATCGGGCACAAAGTTACGGCATTTCGAAGGAAATTAGGGGCTGGAAAG
GGTATAAATACAAAATAGCGCAATAAGTGAAT
```

**References (Supplementary Material)**


**Cloix C, Tutois S, Mathieu O, Cuvillier C, Espagnol MC, Picard G, Tourmente S. 2000**. Analysis of 5S rDNA arrays in *Arabidopsis thaliana*: physical mapping and chromosome-specific polymorphisms. *Genome Res.,* **10**: 679-690.

**Jarvis DE, Ho YS, Lightfoot DJ, Schmöckel SM, Li B, Borm TJA, Ohyanagi H, Mineta K, Michell CT, Saber N, Kharbatia NM, Rupper RR, Sharp AR, Dally N, Boughton BA, Woo YH, Gao G, Schijlen EGWM, Guo X, Momin AA, Negrão S, Al-Babili S, Gehring C, Roessner U, Jung C, Murphy K, Arold ST, Gojobori T, Linden CGvd, van Loo EN, Jellen EN, Maughan PJ, Tester M. 2017**. The genome of *Chenopodium quinoa*. *Nature,* **542**: 307-312.

**Neumann P, Navrátilová A, Koblížková A, Kejnovsky E, Hribova E, Hobza R, Widmer A, Dolezel J, Macas J. 2011**. Plant centromeric retrotransposons: A structural and cytogenetic perspective. *Mob. DNA,* **2**: 4.

**Novák P, Neumann P, Macas J. 2010**. Graph-based clustering and characterization of repetitive sequences in next-generation sequencing data. *BMC Bioinformatics,* **11**: 378.

**Rhoads A, Au KF. 2015**. PacBio sequencing and its applications. *Genomics, Proteomics & Bioinformatics,* **13**: 278-289.

**Ruiz-Ruano FJ, López-León MD, Cabrero J, Camacho JPM. 2016**. High-throughput analysis of the satellitome illuminates satellite DNA evolution. *Scientific Reports,* **6**: 28333.

**Weber B, Heitkam T, Holtgräwe D, Weisshaar B, Minoche AE, Dohm JC, Himmelbauer H, Schmidt T. 2013**. Highly diverse chromoviruses of *Beta vulgaris* are classified by chromodomains and chromosomal integration. *Mob. DNA,* **4**: 8.

**Wheeler TJ, Eddy SR. 2013**. nhmmer: DNA homology search with profile HMMs. *Bioinformatics,* **29**: 2487-2489.
